# Midkine in chick and mouse retinas: neuroprotection, glial reactivity and the formation of Müller glia-derived progenitor cells

**DOI:** 10.1101/2020.08.12.248245

**Authors:** Warren A. Campbell, Amanda Fritsch-Kelleher, Isabella Palazzo, Thanh Hoang, Seth Blackshaw, Andy J. Fischer

**Affiliations:** Department of Neuroscience, College of Medicine, The Ohio State University, Columbus, OH; Solomon H. Snyder Department of Neuroscience, Johns Hopkins University School of Medicine, Baltimore, MD

## Abstract

Recent studies have shown that midkine (MDK), a basic heparin-binding growth factor, is involved in the development and regeneration of the zebrafish retina. However, very little is known about MDK in the retinas of warm-blooded vertebrates. We investigate the expression patterns of MDK and related factors, roles in neuronal survival, and influence upon the formation of Müller glia-derived progenitor cells (MGPCs) in chick and mouse model systems. By using single-cell RNA-sequencing, we find that *MDK* is upregulated during Müller glia (MG) maturation in chick development and when stimulated to reprogram into MGPCs after NMDA damage or FGF2/Insulin treatment. Interestingly, *MDK* is significantly up-regulated by MG in damaged chick retinas, but down-regulated by MG in damaged mouse retinas. In both chick and mouse retinas, exogenous MDK selectively up-regulates cFOS and pS6 (a readout of mTOR-signaling) in MG. In the chick, intraocular injections of MDK before injury is neuroprotective with an observed decrease in dying neurons and microglial reactivity, inducing fewer proliferating MGPCs. Blocking MDK signaling with Na_3_VO_4_ following blocks neuroprotective effects with an increase the number of dying cells and negates the pro-proliferative effects on MGPCs. Inhibitors of PP2A and Pak1 associated with MDK integrin β1 signaling had MG specific inhibitory effects on MGPC formation. In mice, MDK administration with NMDA damage drives a small but significant increase in MGPCs. We conclude that *MDK* expression is dynamically regulated in reactive Müller glia and during reprogramming into MGPCs. MDK acts to coordinate glial activity, neuronal survival, and may act in an autocrine manner to influence the re-programming of Müller glia into proliferating MGPCs.

## Introduction

Midkine (MDK) and pleiotrophin (PTN) are secreted factors that belong to a family of basic heparin-binding cytokines (Muramatsu, 2002). The C-terminal domain of MDK interacts with carbohydrate-bindings proteins which facilitate dimerization and cell signaling (Fabri et al., 1993; Iwasaki et al., 1997; Kilpeläinen et al., 2000; Tsutsui et al., 1991). Extracellular matrix proteoglycans that have a high binding-affinity for MDK include protein tyrosine phosphatase-ζ receptor-like 1 (PTPRZ1), syndecans, glypican-2, PG-M/versican, integrin α_6_β_1_, low density lipoprotein receptor-related protein (LRP), and neuroglycans (Ichihara-Tanaka et al., 2006; Kojima et al., 1996; Kurosawa et al., 2001; Maeda et al., 1999; Mitsiadis et al., 1995; Muramatsu et al., 2000; Muramatsu et al., 2004; Nakanishi et al., 1997; Zou et al., 2000). MDK forms a complex with these proteoglycans to initiate cell-signaling through receptor tyrosine kinases and activation of second messengers such as src, PI-3K, and PAK1 (Qi et al., 2001; Shen et al., 2015; Thillai et al., 2016).

During development the roles of MDK are conserved across many vertebrate species including fish, mice, and humans (Tsutsui et al., 1991). MDK has different functions including promoting cell survival and the proliferation of stem cells, acting directly on stem cells during normal fetal development and organogenesis (Mitsiadis et al., 1995). MDK has been implicated in the pathogenesis of more than 20 different types of cancers, resistance to chemotherapeutics, increased survival of cancerous cells with acidosis and hypoxia, and elevated levels of MDK have been correlated with poor prognoses (Dai et al., 2009; Kang et al., 2004; Mashima et al., 2009; Mirkin et al., 2005; Reynolds et al., 2004; Salama et al., 2006; Takei et al., 2001; Takei et al., 2006; Tsutsui et al., 1993). In damaged mammalian CNS, MDK expression is elevated and may support neuronal survival (Jochheim-Richter et al., 2006; Kikuchi-Horie et al., 2004; Miyashiro et al., 1998; Obama et al., 1998; Sakakima et al., 2006). In rodent eyes, subretinal delivery of MDK protects photoreceptors from light-mediated degeneration (Unoki et al., 1994). In sum, MDK has pleotropic functions that are context-dependent.

In fish, retinal regeneration is a robust process that restores neurons and visual function following damage, whereas this process is far less robust in birds and nearly absent in mammals (Hitchcock and Raymond, 1992; Karl et al., 2008; Raymond, 1991). Müller glia (MG) have been identified as the cell-of-origin for progenitors in mature retinas (Bernardos et al., 2007; Fausett and Goldman, 2006; Fausett et al., 2008; Fischer and Reh, 2001; Ooto et al., 2004). In mammalian retina, significant stimulation, such as forced expression of Ascl1, inhibition of histone deacetylases and neuronal damage, is required to reprogram MG into progenitor-like cells (Karl et al., 2008; Pollak et al., 2013, 1; Ueki et al., 2015). In the chick retina, MG readily reprogram into progenitor-like cells that proliferate, but the progeny have a limited capacity to differentiate as neurons (Fischer and Reh, 2001; Fischer and Reh, 2003). Understanding the mechanisms that regulate the proliferation and differentiation of MGPCs is important to harnessing the regenerative potential of MG in warm-blooded vertebrates.

Recent studies in zebrafish retina have indicated that MDK-a and MDK-b are up-regulated in stem niches and in MG during reprogramming (Calinescu et al., 2009). MDK-a is expressed by mitotic retinal progenitors at 30 hrs post-fertilization, then in MG at 72 hrs post-fertilization through adulthood (Gramage et al., 2014). Knock-down of MDK-b results in microphthalmia or anophthalmia (Calinescu et al., 2009). During reprogramming of MG into MGPCs following retinal damage, MDK-a controls cell cycle exit and neuronal differentiation via bHLH transcription factor Id2a (Luo et al., 2012; Nagashima et al., 2019). Nothing is known about how MDK influences the process of retinal regeneration in warm-blooded vertebrates. Accordingly, we investigate expression patterns and the impact of MDK on glial cells in the chick and mouse retinas *in vivo.*

## Methods and Materials

### Animals

The animals approved for use in these experiments was in accordance with the guidelines established by the National Institutes of Health and IACUC at The Ohio State University. Newly hatched P0 wildtype leghorn chicks (*Gallus gallus domesticus*) were obtained from Meyer Hatchery (Polk, Ohio). Post-hatch chicks were maintained in a regular diurnal cycle of 12 hours light, 12 hours dark (8:00 AM-8:00 PM). Chicks were housed in stainless-steel brooders at 25°C and received water and Purina^tm^ chick starter *ad libitum.* Mice were kept on a cycle of 12 h light, 12 h dark (lights on at 6:00 AM). C57BL/6J mice between the ages of P60-P100 were used for all experiments.

Fertilized eggs were obtained from the Michigan State University, Department of Animal Science. Eggs were incubated at a constant 37.5°C, with a 1hr period room temperature cool down every 24hrs. Additionally, the eggs were rocked every 45 minutes, and held at a constant relative humidity of 45%. Embryos were harvested at various time points after incubation and staged according to guidelines established by Hamburger and Hamilton (1951).

### Intraocular injections

Chicks were anesthetized with 2.5% isoflurane mixed with oxygen from a non-rebreathing vaporizer. The technical procedures for intraocular injections were performed as previously described (Fischer et al., 1998). With all injection paradigms, both pharmacological and vehicle treatments were administered to the right and left eye respectively. Compounds were injected in 20 μl sterile saline with 0.05 mg/ml bovine serum albumin added as a carrier. For mice injections, the total volume injected into each eye was 2μl. The details of compounds injected in to the vitreous are described (Table S1)

### Preparation of clodronate liposomes

Clodronate liposomes were synthesized utilizing a modified protocol from previous descriptions (Van Rooijen, 1989; van Rooijen, 1992; Zelinka et al., 2012). In short, approximately 8 mg of L-α-Phosphatidyl-DL-glycerol sodium salt (Sigma P8318) was dissolved in chloroform. 50 mg of cholesterol was dissolved in chloroform with the lipids in a sterile microcentrifuge tube. This tube was rotated under nitrogen gas to evaporate the chloroform and leave a thin lipid-film on the walls of the tube. 158 mg dichloro-methylene diphosphonate (clodronate; Sigma-Aldrich) dissolved sterile PBS (pH 7.4) was added to the lipid/cholesterol film and vortexed for 5 minutes. To reduce size variability of lipid vesicles, the mixture was sonicated at 42 kHz for 6 minutes. Purification of liposomes was accomplished via centrifugation at 10,000 x G for 15 minutes, aspirated, and resuspended in 150 μl PBS. Each retinal injection used between 5 and 20 ul of clodronate-liposome solution. There was a variable yield of clodronate-liposomes during the purification resulting in some variability per dose. The dosage was adjusted such that >98% of the microglia are ablated by 2 days after administration with no off-target cell death or pigmented epithelial cells.

### Single Cell RNA sequencing of retinas

Retinas were obtained from embryonic, postnatal chick, and adult mouse retinas. Isolated retinas were dissociated in a 0.25% papain solution in Hank’s balanced salt solution (HBSS), pH = 7.4, for 30 minutes, and suspensions were frequently triturated. The dissociated cells were passed through a sterile 70μm filter to remove large particulate debris. Dissociated cells were assessed for viability (Countess II; Invitrogen) and cell-density diluted to 700 cell/μl. Each single cell cDNA library was prepared for a target of 10,000 cells per sample. The cell suspension and Chromium Single Cell 3’ V2 reagents (10X Genomics) were loaded onto chips to capture individual cells with individual gel beads in emulsion (GEMs) using 10X Chromium Controller. cDNA and library amplification for an optimal signal was 12 and 10 cycles respectively. Sequencing was conducted on Illumina HiSeq2500 (Genomics Resource Core Facility, John’s Hopkins University) or HiSeq4000 (Novogene) with 26 bp for Read 1 and 98 bp for Read 2. Fasta sequence files were de-multiplexed, aligned, and annotated using the chick ENSMBL database (GRCg6a, Ensembl release 94) or mouse ENSMBL database (GRCm38.p6, Ensembl release 67) and Cell Ranger software. Gene expression was counted using unique molecular identifier bar codes, and gene-cell matrices were constructed. Using Seurat toolkits, t-distributed stochastic neighbor embedding (tSNE) plots or Uniform Manifold Approximation and Projection for Dimension Reduction (UMAP) plots were generated from aggregates of multiple scRNA-seq libraries (Butler et al., 2018; Satija et al., 2015). Compiled in each tSNE/UMAP plot are two biological library replicates for each experimental condition. Seurat was used to construct violin/scatter plots. Significance of difference in violin/scatter plots was determined using a Wilcoxon Rank Sum test with Bonferroni correction. Monocle was used to construct unbiased pseudo-time trajectories and scatter plotters for MG and MGPCs across pseudotime (Qiu et al., 2017a; Qiu et al., 2017b; Trapnell et al., 2012). Genes that were used to identify different types of retinal cells included the following: (1) Müller glia: *GLUL, VIM, SCL1A3, RLBP1,* (2) MGPCs: *PCNA, CDK1, TOP2A, ASCL1,* (3) microglia: *C1QA, C1QB, CCL4, CSF1R, TMEM22,* (4) ganglion cells: *THY1, POU4F2, RBPMS2, NEFL, NEFM,* (5) amacrine cells: *GAD67, CALB2, TFAP2A,* (6) horizontal cells: *PROX1, CALB2, NTRK1,* (7) bipolar cells: *VSX1, OTX2, GRIK1, GABRA1*, and (7) cone photoreceptors: *CALB1, GNAT2, OPN1LW,* and (8) rod photoreceptors: *RHO, NR2E3, ARR3.* The MG have an over-abundant representation in the scRNA-seq databases. This likely resulted from fortuitous capture-bias and/or tolerance of the MG to the dissociation process. Single cell libraries for NMDA and FGF insulin in the mouse and chick generated by our lab have first been reported for cross species comparisons (Hoang et al., 2019), where chick embryonic retinal libraries are first being reported in this study.

### Fixation, sectioning and immunocytochemistry

Retinal tissue samples were formaldehyde fixed, sectioned, and labeled via immunohistochemistry as described previously (Fischer et al., 2008; Fischer et al., 2009a). Antibody dilutions and commercial sources for images used in this study are described (Table S2). Observed labeling was not due to off-target labeling of secondary antibodies or tissue autofluorescence because sections incubated exclusively with secondary antibodies were devoid of fluorescence. Secondary antibodies utilized include donkey-anti-goat-Alexa488/568, goat-anti-rabbit-Alexa488/568/647, goat-anti-mouse-Alexa488/568/647, goat-anti-rat-Alexa488 (Life Technologies) diluted to 1:1000 in PBS and 0.2% Triton X-100.

### Labeling for EdU

For the detection of nuclei that incorporated EdU, immunolabeled sections were fixed in 4% formaldehyde in 0.1M PBS pH 7.4 for 5 minutes at room temperature. Samples were washed for 5 minutes with PBS, permeabilized with 0.5% Triton X-100 in PBS for 1 minute at room temperature and washed twice for 5 minutes in PBS. Sections were incubated for 30 minutes at room temperature in a buffer consisting of 100 mM Tris, 8 mM CuSO4, and 100 mM ascorbic acid in dH2O. The Alexa Fluor 568 Azide (Thermo Fisher Scientific) was added to the buffer at a 1:100 dilution.

### Terminal deoxynucleotidyl transferase dUTP nick end labeling (TUNEL)

The TUNEL assay was implemented to identify dying cells by imaging fluorescent labeling of double stranded DNA breaks in nuclei. The *In Situ* Cell Death Kit (TMR red; Roche Applied Science) was applied to fixed retinal sections as per the manufacturer’s instructions.

### Photography, measurements, cell counts and statistics

Microscopy images of retinal sections were captured with the Leica DM5000B microscope with epifluorescence and the Leica DC500 digital camera. High resolution confocal images were obtained with a Leica SP8 available in The Department of Neuroscience Imaging Facility at The Ohio State University. Representative images are modified to have enhanced color, brightness, and contrast for improved clarity using Adobe Photoshop. In EdU proliferation assays, a fixed region of retina was counted and average numbers of Sox2 and EdU co-labeled cells. The retinal region selected for investigation was standardized between treatment and control groups to reduce variability and improve reproducibility.

Similar to previous reports (Fischer et al., 2009a; Fischer et al., 2009b; Ghai et al., 2009), immunofluorescence was quantified by using ImagePro6.2 (Media Cybernetics, Bethesda, MD, USA) or Image J (NIH). Identical illumination, microscope, and camera settings were used to obtain images for quantification. Retinal areas were sampled from 5.4 MP digital images. These areas were randomly sampled over the inner nuclear layer (INL) where the nuclei of the bipolar and amacrine neurons were observed. Measurements of immunofluorescence were performed using ImagePro 6.2 as described previously (Ghai et al., 2009; Stanke et al., 2010; Todd and Fischer, 2015). The density sum was calculated as the total of pixel values for all pixels within thresholded regions. The mean density sum was calculated for the pixels within threshold regions from ≥5 retinas for each experimental condition. GraphPad Prism 6 was used for statistical analyses.

Measurement for immunofluorescence of cFos in the nuclei of MG/MGPCs were made by from single optical confocal sections by selecting the total area of pixel values above threshold (≥70) for Sox2 or Sox9 immunofluorescence (in the red channel) and copying nuclear cFos from only MG (in the green channel). The MG-specific cFos was quantified (as described below or copied onto a 70% grayscale background for figures. Measurements of pS6 immunofluorescence were made for a fixed, cropped area (14,000 μm^2^) of INL, OPL and ONL. Measurements were made for regions containing pixels with intensity values of 70 or greater (0 = black and 255 = saturated). The total area was calculated for regions with pixel intensities above threshold. The intensity sum was calculated as the total of pixel values for all pixels within threshold regions. The mean intensity sum was calculated for the pixels within threshold regions from ≥5 retinas for each experimental condition.

For statistical evaluation of differences in treatments, a two-tailed paired *t*-test was applied for intra-individual variability where each biological sample also served as its own control. For two treatment groups comparing inter-individual variability, a two-tailed unpaired *t*-test was applied. For multivariate analysis, an ANOVA with the associated Tukey Test was used to evaluate any significant differences between multiple groups.

## Results

### *MDK* and *PTN* in embryonic retina

scRNA-seq retina libraries were established at four stages of development including E5, E8, E12, and E15. The aggregation of these libraries yielded 22,698 cells after filtering to exclude doublets, cells with low UMI, and low genes/cell. UMAP plots of aggregate libraries of embryonic retinas formed clustered of cells into patterns that correlated to both developmental stage and cell type (Fig. 1a). Cell types were identified based on expression of well-established markers. Specifically, retinal progenitor cells from E5 and E8 retinas were identified by expression of *ASCL1, CDK1,* and *TOP2A.* (Supplemental Fig. 1a,b). Maturing MG were identified by expression of *GLUL, RLBP1* and *SLC1A3* (Supplemental Fig. 1a,b).

**Figure 1.**
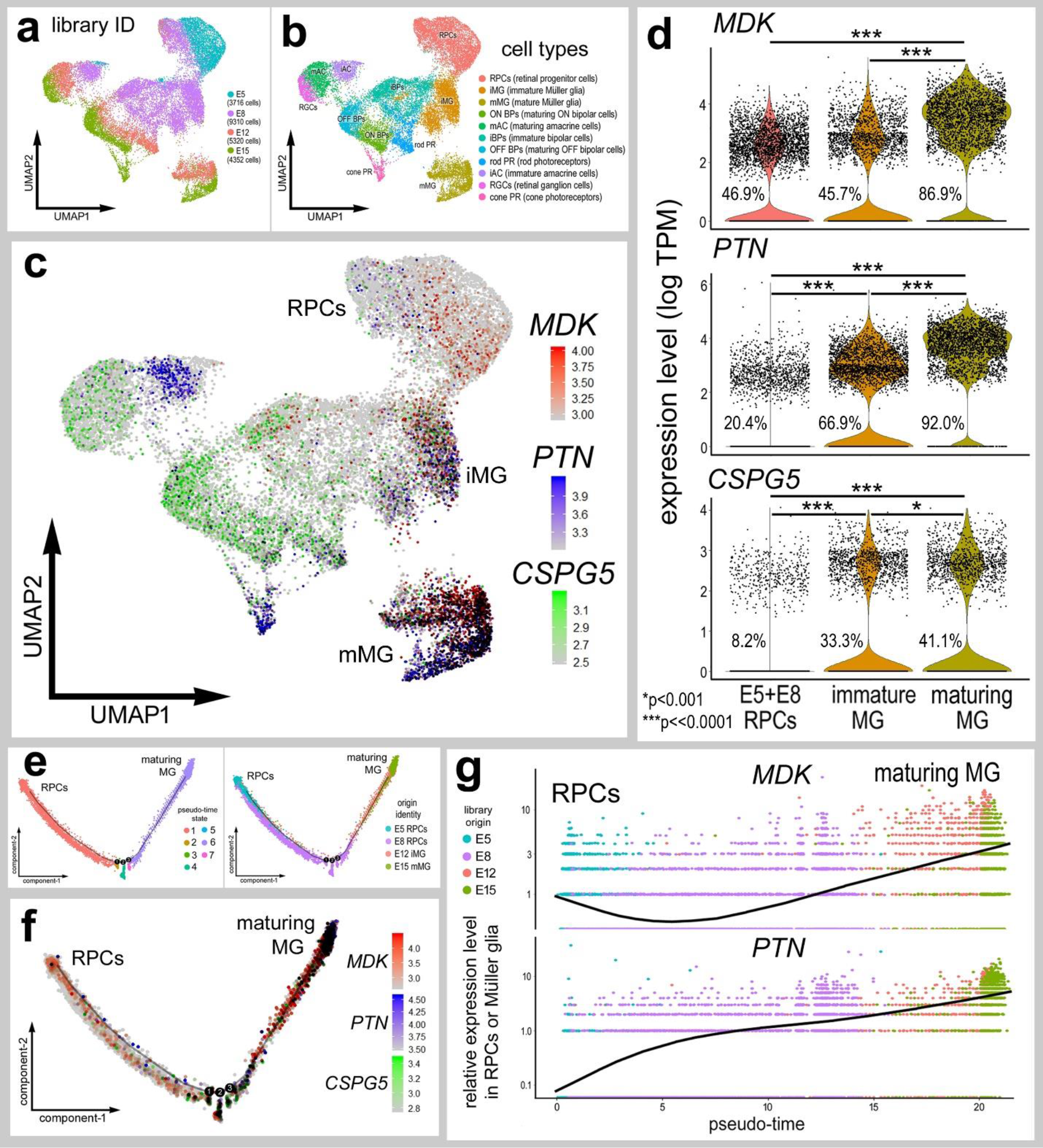
Expression of *MDK, PTN* and *CSPG5* in maturing MG in embryonic chick retina. scRNA-seq was used to identify patterns of expression of *MDK, PTN* and putative receptor *CSPG5* among embryonic retinal cells at four stages of development (E5, E8, E12, E15). UMAP-ordered clusters of cells were identified by expression of hallmark genes (**a,b**). A heatmap of *MDK, PTN* and *CSPG5* illustrates expression profiles in different developing retinal cells (**c**). Each dot represents one cell and black dots indicate cells with 2 or more genes expressed. The upregulation of MDK and PTN in RPCs and maturing MG is illustrated with violin plot (**d**). The number on each violin indicates the percentage of expressing cells. The transition from RPC to mature MG is modelled with pseudotime ordering of cells with early RPCs to the far left and maturing MG to the right of the pseudotime trajectory (**e**). *MDK* and *PTN* are up-regulated in MG during maturation as illustrated by the pseudotime heatmap (**f**) and pseudotime plot (**g**). Significant difference (*p<0.01, **p<0.0001, ***p<<0.0001) was determined by using a Wilcox rank sum with Bonferoni correction. RPC – retinal progenitor cell, MG – Müller glia, iMG – immature Müller glia, mMG - mature Müller glia.

Elevated levels of *MDK* expression were observed in MG at E12 and E15, with lower levels of expression in immature MG at E8 and retinal progenitor cells at E5 (Fig. 1c,d). *PTN* was prominently expressed in immature and mature MG, but was also detected in immature amacrine cells (E8), rod photoreceptors (E12), and cone photoreceptors (E15) (Fig. 1c). Levels of *MDK* and *PTN* were significantly higher in maturing MG compared to immature MG and RPCs (Fig. 1d). Putative receptors and signal transducers of MDK and PTN include integrin β1 (*ITGB1),* receptor-like protein tyrosine phosphatase-ζ (*PTPRZ1),* chondroitin sulfate proteoglycan 5 (*CSPG5*) and p21-activated serine/threonine kinase (*PAK1).* These mRNAs had variable, scattered expression in embryonic retinal cells. Scattered expression of *PTPRZ1, ITGB1* and *PAK1* was observed in RPCs, immature and mature MG (Supplemental Fig. 1c). *CSPG5* was expressed by developing photoreceptors, amacrine, ganglion and bipolar cells (Fig. 1c). Additionally, *CSPG5* was expressed at significantly elevated levels by immature and maturing MG compared to levels in RPCs (Fig. 1c.d).

Re-embedding of RPCs and MG for pseudotime analysis revealed a trajectory of cells with early RPCs and maturing MG at opposite ends of the trajectory (Fig. 1e). Across the pseudotime trajectory levels of *GLUL* increased, while levels of *CDK1* decreased (Supplemental Fig. 1e,f). Similar to the pattern of expression of *GLUL,* the expression of *MDK* and *PTN* increases from retinal progenitors to maturing MG (Fig. 1e,f). Higher levels of expression were observed for both *MDK* and *PTN*, with *PTN* at low levels in retinal progenitors (Fig. 1f,g). Across pseudotime, levels of *MDK* were high in early progenitors, with a dip in expression during transition phases, and increased in maturing MG (Fig. 1f,g). Collectively, these findings suggest that both *PTN* and *MDK* are up-regulated by maturing MG during late stages of embryonic development, and, based on patterns of expression of putative receptors, MDK and PTN may have autocrine and paracrine actions in late-stage embryonic chick retina.

### Upregulation MDK in MG of damaged retinas

scRNA-seq libraries were aggregated for retinal cells obtained from control and NMDA-damaged retinas at various time points (24, 48 and 72 hrs) after treatment (Fig. 2a). UMAP plots were generated and clusters of different cells were identified based on well-established patterns of expression (Fig. 2a,b). For example, resting MG formed a discrete cluster of cells and expressed high levels of *GLUL, RLBP1* and *SLC1A3* (Supplemental Fig. 2a,b). After damage, MG down-regulate markers of mature glia as they transition into reactive glial cells and into progenitor-like cells that up-regulate *TOP2A, CDK1* and *ESPL1* (Supplemental Fig. 2a,b). *MDK* was expressed at low levels in relatively few resting MG in undamaged retina, unlike maturing MG in late stages of embryonic retinas (Fig. 1c,e), suggesting a down-regulation of *MDK* in MG as development proceeds after hatching. The expression of *MDK* was scattered in oligodendrocytes and Non-astrocytic Inner Retinal Glia (NIRGs). NIRG cells are a distinct type of glial cells that has been described in the retinas of birds (Rompani and Cepko 2010; Fischer et al., 2010) and some types of reptiles (Todd et al., 2016). Following NMDA-induced damage, *MDK* is upregulated in MG and MGPCs at 24hrs, 48hrs, and 72hrs after treatment (Fig. 2c,e). In addition, MGPCs maintain high levels of *MDK* (Fig. 2c,e). By comparison, *PTN* was widely expressed in most types of retinal cells, and was significantly down-regulated by MG and MGPCs in damaged retinas (Fig. 2c,e). We queried expression of putative receptors for *MDK* and *PTN*, including *PTPRZ1, CSPG5* and Syndecan 4 (*SDC4).* Although *SDC4* was expressed in scattered retina cells, *SDC4* was low in resting MG and was up-regulated across MG at 24hrs after NMDA-treatment (Fig. 2d,e). *CSPG5* was expressed at high levels in resting MG, and was down-regulated in activated MG at 24hrs after NMDA and in MGPCs, and remained elevated in activated MG at 48 and 72hrs after NMDA (Fig. 2d,e). *PTPRZ1* was not detected in MG, but was expressed at high levels in NIRG cells and in scattered amacrine and bipolar cells (Fig. 2d). *CSPG5, SDC4, ITGB1,* and *PTN* were dynamically expressed by different retinal neurons and NIRG cells in NMDA-damaged retinas (Supplemental Fig. 2a-e), suggesting that MDK may influence NIRG cells and neurons following an insult.

**Figure 2.**
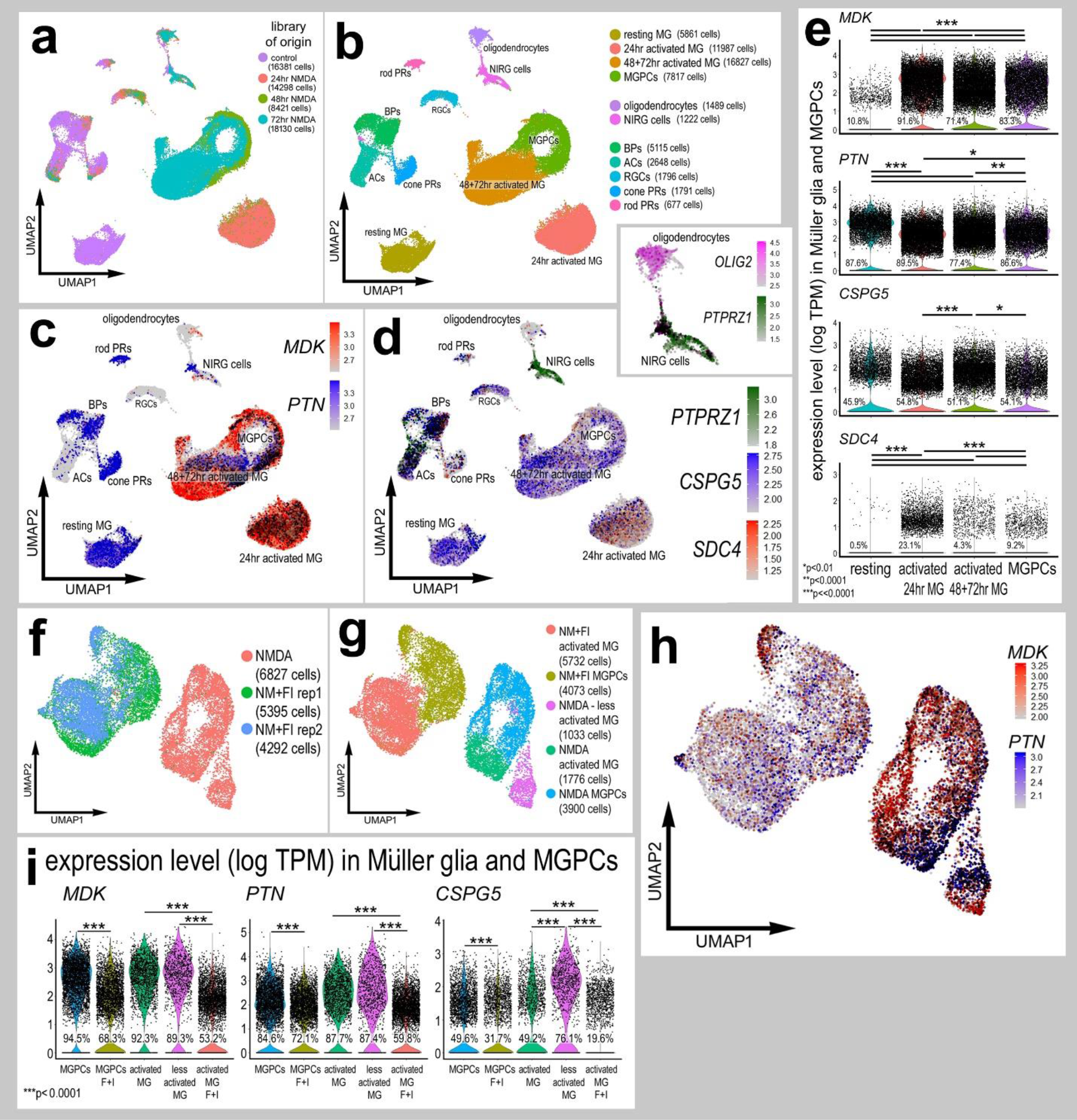
The expression profile of *MDK, PTN* and putative receptors in mature retinal cells and following acute injury. scRNA-seq was used to identify patterns of expression of MDK-related genes among acutely dissociated retinal cells with the data presented in UMAP plots (**a**-**d, f, g, h**) and violin plots (**e**,**i**). Control and treated scRNA-seq libraries were aggregated from 24hr, 48hr, and 72hr after NMDA-treatment (**a**). UMAP-ordered cells formed distinct clusters with MG and MGPCs forming distinct clusters (**b**). Expression heatmaps of *MDK, PTN,* and receptor genes *PTPRZ1, CSPG5,* and *SDC4* demonstrate patterns of expression in the retina, with black dots representing cells with 2 or more genes (**c,d**). In addition to NMDA, retinas were treated with insulin and FGF2 and expression levels of *MDK, PTN,* and *CSPG5* were assessed in MG and MGPCs (**f-i**). UMAP and violin plots illustrate relative levels of expression in MG and MGPCs treated with NMDA alone or NMDA plus insulin and FGF2 (**h,i**). Violin plots illustrate levels of gene expression and significant changes (*p<0.1, **p<0.0001, ***p<<0.0001) in levels were determined by using a Wilcox rank sum with Bonferroni correction. The number on each violin indicates the percentage of expressing cells.

Analysis of MG and MGPCs in different pseudotime states revealed a branched trajectory with resting MG, proliferating MGPCs, and activated MG from 72hr after NMDA-treatment largely confined to different branches and states (Supplemental Fig. 3a-d). The expression of *MDK* across pseudotime positively correlates with a transition toward an MGPC-phenotype and up-regulation of progenitor markers, such as *CDK1,* and inversely correlated to resting glial phenotypes with significant down-regulation of glial markers such as *GLUL* (Supplemental Fig. 3a-d). By comparison, levels of *PTN* were decreased across pseudotime, with the largest decrease in *PTN* in activated MG compared to resting MG (Supplemental Fig. 3a-d). Similar to expression patterns of *CDK1,* patterns of expression of *SDC4* are significantly elevated in pseudotime state 4 populated by activated MG and proliferating MGPCs (Supplemental Fig. 3a-d).

When comparing MG and MGPCs from 48hrs after NMDA with and without FGF2 and insulin, the relative levels of *MDK* and *PTN* were significantly decreased (Fig. 2f-i). UMAP plots revealed distinct clustering of MG and MGPCs from retinas from 48hrs NMDA alone and 48hrs NMDA plus FGF2 and insulin (Fig. 2f-i). Similarly, levels of *GLUL, RLBP1* and *CSPG5* were significantly decreased by FGF2 and insulin in damaged retinas in both MG and MGPCs (Fig. 2f-I; Supplemental Fig. 3e-g). By contrast, levels of *CDK1* and *TOP2A* were significantly increased by FGF2 and insulin in MGPCs in damaged retinas (Supplemental Fig. 3e-g). Collectively, these findings suggest expression levels of *PTN* reflects a resting glial phenotype and levels are decreased by damage and further decreased by FGF2 and insulin, whereas expression levels of *MDK* corresponds with acutely activated glia or maturing glia, which is strongly induced by damage, but decreased by FGF2 and insulin in damaged retinas.

### MDK, neuroprotection and glial reactivity in damaged retinas

The large, significant up-regulation of *MDK* by MG in NMDA-treated retinas suggests that this growth factor is involved in the responses of retinal cells to acute damage. To determine whether MDK influences retinal cells we probed for the activation of different cell-signaling pathways following a single intraocular injection of recombinant MDK or PTN. Four hours after delivery of MDK we found a significant up-regulation of cFOS and pS6 specifically in MG (Fig. 3a-e), suggesting activation of the mTor-pathway. In addition, amacrine cells appeared to significantly up-regulate cFOS and NIRG cells up-regulated pS6 in response to MDK (Fig. 3a-e). To test whether cell signaling was influenced by PP2A-inhibitors, we co-applied fostriecin and calyculin A with MDK. We found that fostriecin and calyculin A significantly reduced levels of pS6 in MDK-treated MG (Fig. 3f,g), where cfos activation was unaffected and independent of PP2A inhibition (data not shown). We failed to find up-regulation of pERK1/2, p38 MAPK, pCREB, pSmad1/5/8, pStat3, and nuclear smad2/3 following intravitreal delivery of MDK (not shown). We failed to detect changes in cell signaling in response to intraocular injections of PTN (not shown). Despite distinct activation of cFOS and mTor in MG from a single dose of MDK, four consecutive daily intraocular injections of MDK or PTN had no significant effect upon MG reactivity or formation of proliferating MGPCs (not shown).

**Figure 3.**
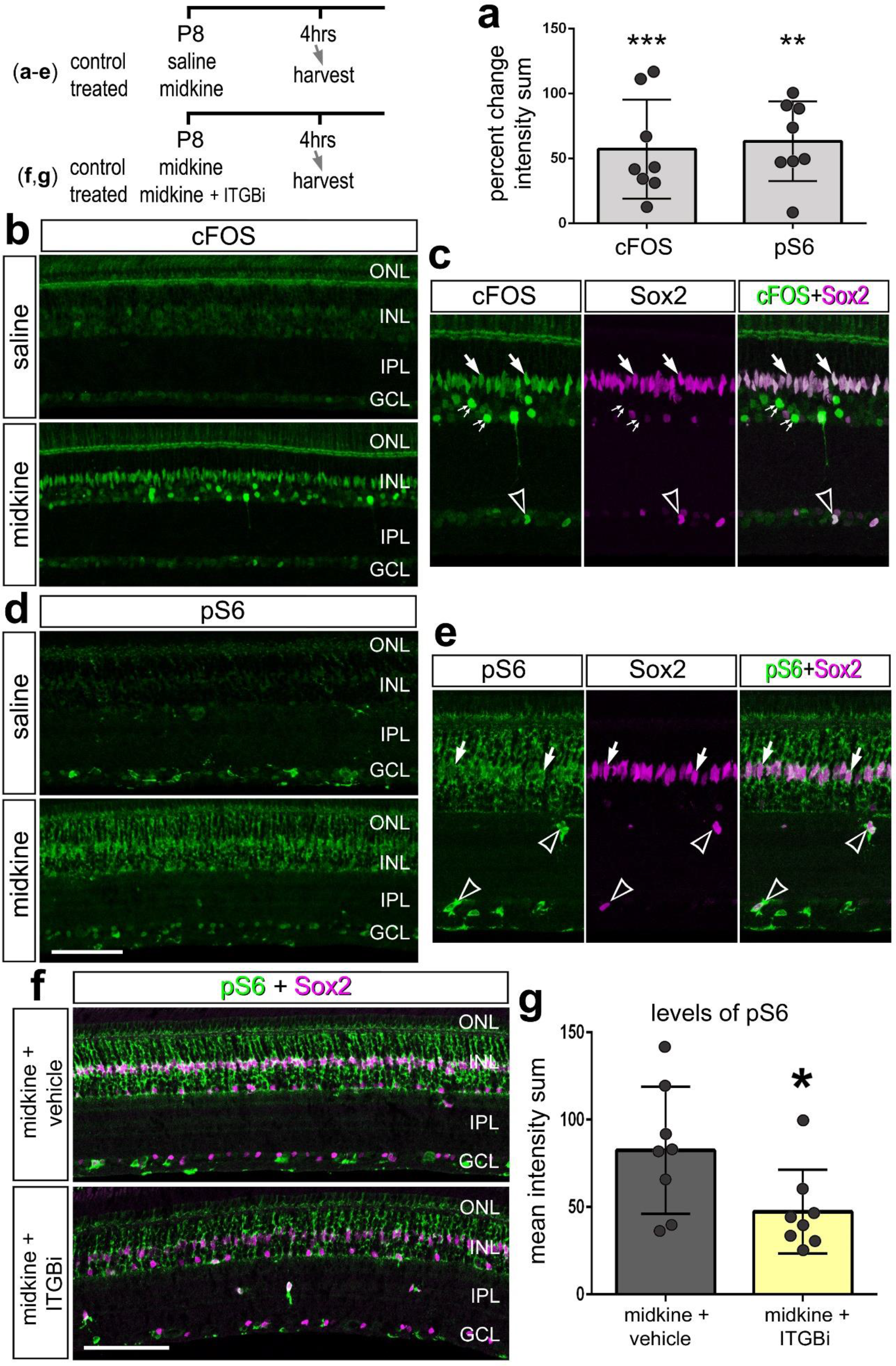
MDK activates cell-signaling in MG in the chick retina. A single intraocular injection of MDK was delivered and retinas were harvested 4 hours later. The histogram in **a** represents the mean percent change (±SD) in intensity sum for cFOS and pS6 immunofluorescence. Each dot represents one biological replicate retina. Significance of difference (**p<0.01, ***p<0.001) was determined by using a paired *t*-test. Sections of saline (control) and MDK-treated retinas were labeled with antibodies to cFOS (green; **b**,**c**), pS6 (green; **d**,**e**) and Sox2 (magenta; **c**,**e**). Arrows indicate the nuclei of MG, small double-arrows indicate the nuclei of amacrine cells, and hollow arrow-heads indicate the nuclei of presumptive NIRG cells. An identical paradigm with the addition of ITGB1 inhibitors fostriecin & calyculin measured changes in pS6 signaling in MG (**f**) and was quantified for intensity changes (**g**). The calibration bar (50 μm) in panel **d** applies to **b** and **d**. Abbreviations: ONL – outer nuclear layer, INL – inner nuclear layer, IPL – inner plexiform layer, GCL – ganglion cell layer.

Intravitreal injections of MDK after NMDA-treatment had no significant effect upon glial reactivity or proliferation of MGPCs, NIRG cells or microglia (not shown). Concurrent administration lacked significant effects because endogenous levels of MDK were very high and MDK-mediated cell-signaling may have been saturated. By comparison, injection of MDK prior to NMDA-treatment significantly reduced the numbers of proliferating MGPCs that accumulated EdU or were immunolabeled for pHisH3 (Fig. 4a-d). Similarly, there was a significant reduction in the number Sox2^+^/Nkx2.2^+^ NIRG cells that accumulated in NMDA-damaged retinas (Fig. 4e,f). To determine whether microglial reactivity was influenced by MDK we measured the area and intensity sum for CD45 immunofluorescence, which is increased in reactive microglia (Fischer et al., 2014). Both the area and the intensity of CD45-immunofluorescence was decreased in response to MDK (Fig. 4g,h). We failed to detect a significant MDK-mediated change in well-established read-outs of different cell-signaling pathways including pS6, pCREB, p38 MAPK, pERK1/2, or pStat3 at later time points after damage (data not shown).

**Figure 4.**
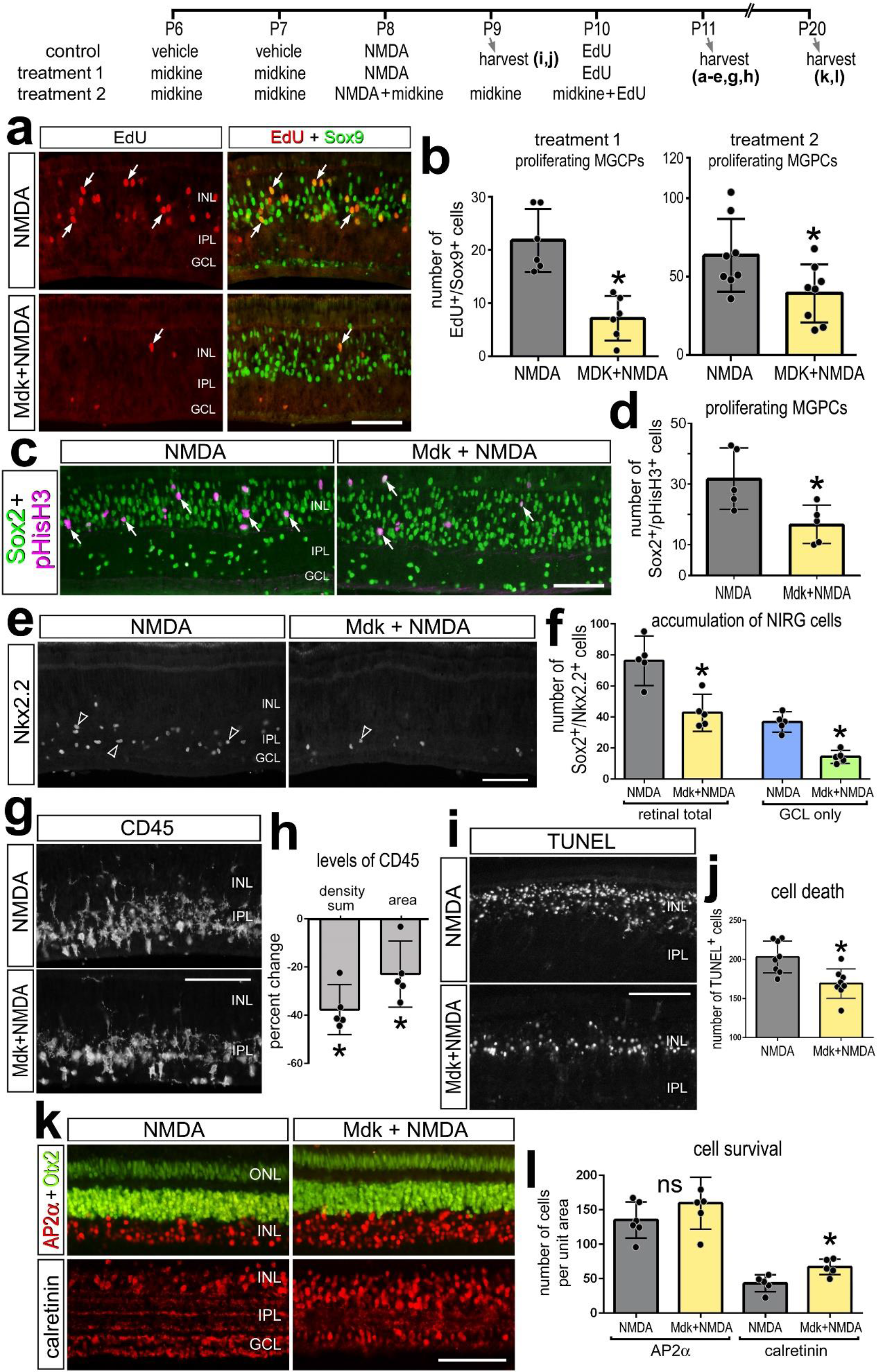
MDK treatment prior to NMDA reduces numbers of proliferating MGPCs, suppresses the accumulation of NIRG cells, and increases neuronal survival. Eyes were injected with MDK or saline at P6 and P7, and NMDA at P8. Some retinas were harvested at P9, whereas other eyes were injected at P10 with EdU and retinas harvested 4hrs later, 24hrs later at P11 or 10 days later at P20. Sections of the retina were labeled for EdU (red) and Sox9 (green; **a**), phospho-Histone H3 (pHisH3; magenta) and Sox2 (green; **c**), Nkx2.2 (**e**), CD45 (**g**), TUNEL (**i**), and AP2α (red) and Otx2 (green) or calretinin (red; **k**). Arrows indicate nuclei of proliferating MGPCs and hollow arrow-heads indicate TUNEL-positive cells. The histogram/scatter-plots **b**, **d, f, j** and **l** illustrate the mean number of labeled cells (±SD). The histogram in **h** represents the mean percent change (±SD) in density sum and area for CD45 immunofluorescence. Each dot represents one biological replicate. Significance of difference (*p<0.01) was determined by using a paired *t*-test. The calibration bars panels **a**, **c, e, g, i** and **k** represent 50 μm. Abbreviations: ONL – outer nuclear layer, INL – inner nuclear layer, IPL – inner plexiform layer, GCL – ganglion cell layer.

Levels of retinal damage and cell death positively correlate to numbers of proliferating MGPCs (Fischer and Reh, 2001; Fischer and Reh, 2003). Thus, it is possible that reduced numbers of proliferating MGPCs resulted from less cell death with MDK pre-treatment. Using the TUNEL method to labeling dying cells, we found that the administration of MDK before NMDA-damage significantly reduced numbers of dying cells (Fig. 4i,j). Decreased numbers of dying cells were observed at both 24h and 72h after NMDA-treatment with MDK pre-treatment. To complement the cell death studies, we probed for long-term survival of inner retinal neurons. Although there was no change in numbers of AP2α^+^ amacrine cells, there was a significant increase in numbers of calretinin^+^ cells in retinas treated with MDK (Fig. 4k,i).

*PTN* was significantly down-regulated in MG following NMDA-treatment (Fig. 2). Despite the administration of high doses (1μg/dose) of PTN, intravitreal delivery of PTN with NMDA had no measurable effects on the formation of MGPCs, the reactivity and proliferation of microglia, and accumulation of NIRG cells (data not shown). Although these experiments were conducted using recombinant human PTN, there is high conservation between chick, mouse and human PTN (92% & 93% respectively).

### Inhibition of MDK-signaling in damaged retinas

MDK-signaling is often upregulated in tissues with proliferating cells, such as tumors, and this proliferation can be suppressed by inhibition of MDK-signaling (Hao et al., 2013; Takei et al., 2006). Since levels of *MDK* were markedly increased in MG in damaged retinas, we tested whether inhibition of MDK-signaling influenced the formation of proliferating MGPCs. We applied a MDK-expression inhibitor (MDKi), which downregulates protein expression in a dose dependent manner (Masui et al., 2016). However, application of MDKi after NMDA-treatment failed to influence MG, NIRG cells, or microglia (data not shown). Alternatively, we applied a PTPRZ inhibitor SCB4380 that targets the intracellular domain of the receptor (Fujikawa et al., 2016). However, this inhibitor failed to influence MG, NIRG cells or microglia when applied after NMDA-treatment (not shown). It is likely that MDKi and SCB4380 had poor cellular permeability or species specificity and these drugs failed to adequately diffuse into the retina and mediate cellular changes.

We next applied an inhibitor of MDK-signaling, sodium orthovanadate (Na_3_VO_4_) that suppresses the activity of tyrosine phosphatase activity including PTPRZ1 (Qi et al., 2001), which was predominantly expressed by NIRG cells, amacrine cells, and some bipolar cells (Fig. 2c). Application of Na_3_VO_4_ with and after NMDA significantly reduced numbers of proliferating MGPCs (Fig. 5a,b). In addition, treatment with Na_3_VO_4_ significantly increased numbers of NIRG cells in the IPL (Fig. 5c,d) and increased numbers of dying cells (Fig. 5e,f). Despite this increase in retinal damage, proliferation of MGPCs was reduced in response to Na_3_VO_4_ after NMDA-induced damage (Fig. 5a,b).

**Figure 5.**
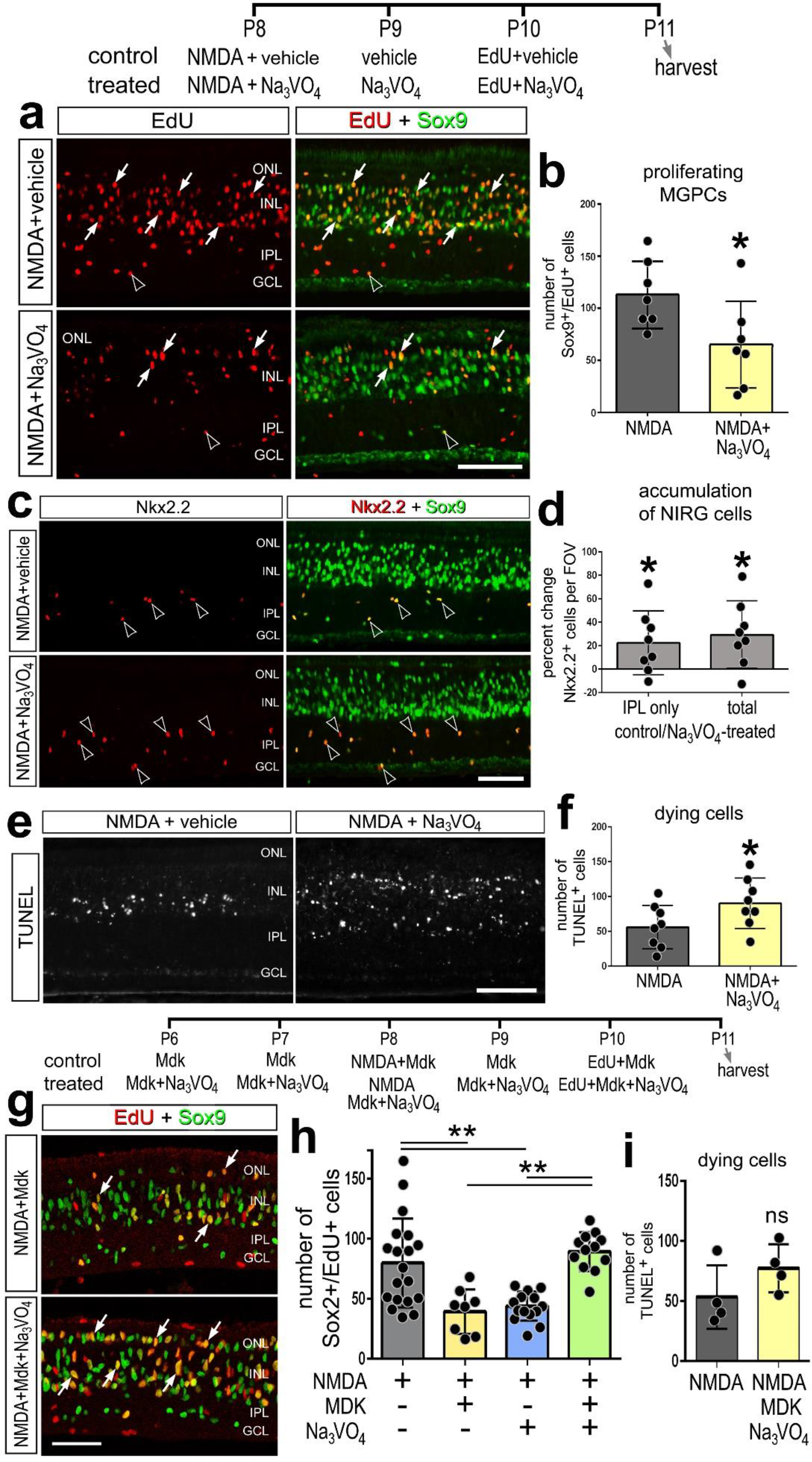
Inhibition of PTPrz in NMDA-damaged retinas suppressed the formation of MGPCs, increases numbers of dying cells, and stimulates the accumulation of NIRG cells. Eyes were injected with NMDA and Na_3_VO_4_ inhibitor or vehicle at P8, inhibitor or vehicle at P9, EdU at P10, and retinas harvested at P11. Sections of the retina were labeled for EdU (red) and Sox9 (green; **a, g**), Nkx2.2 and Sox9 (green; **c**), or TUNEL (**e**). Arrows indicate nuclei of proliferating MGPCs (**a,g**) and hollow arrow-heads indicate NIRG cells (**c**). The histogram/scatter-plots in **b**, **d, f, h** and **i** illustrate the mean (±SD) number of labeled cells. Each dot represents one biological replicate. Significance of difference (*p<0.05) was determined by using a paired *t*-test. The calibration bars panels **a**, **c, e** and **g** represent 50 μm. Abbreviations: ONL – outer nuclear layer, INL – inner nuclear layer, IPL – inner plexiform layer, GCL – ganglion cell layer.

We next examined the specificity of Na_3_VO_4_, by testing whether Na_3_VO_4_ blocked the effects of MDK when applied with NMDA-treatment. Comparison across treatment groups (NMDA alone, NMDA + Na_3_VO_4_, NMDA + MDK, and NMDA + Na_3_VO_4_ + MDK) revealed a significant decrease in MGPCs in eyes treated with Na_3_VO_4_ and MDK alone (Fig. 5g,h). With the combination of MDK and Na_3_VO_4_ there was a significant increase in proliferating MGPCs relative to treatment with MDK or Na_3_VO_4_ alone (Fig. 5h). However, this level was not increased relative to levels seen with NMDA alone (Fig. 5h). In addition, the combination of MDK and Na_3_VO_4_ resulted in no significant difference in numbers of TUNEL^+^ dying cells compared to NMDA alone (Fig 5i). These findings suggest that the effects of MDK and Na_3_VO_4_ upon proliferating MGPCs and numbers of dying neurons are mediated by overlapping cellular targets.

### Putative MDK receptors, signal transducers and MGPC formation

MDK has been found to bind and signal through Integrin-Beta 1 (ITGB1) (Muramatsu et al., 2004). ITGB1 signaling through secondary messengers Integrin linked kinase (ILK), p21 activated kinase 1 (PAK1), cell division factor 42 (CDC42), protein phosphatase 2a (PP2A, gene: *PPP2CA),* and Git/Cat-1 regulate cytoskeleton remodeling, migration, and cellular proliferation (Bagrodia and Cerione, 1999; Ivaska et al., 1999; Kawachi et al., 2001; Kim et al., 2004; Martin et al., 2016; Mulrooney et al., 2000) (see Fig. 12). *PAK1* has been implicated as a cell cycle regulator that is downstream of MDK and PTN signaling (Kawachi et al., 2001). PAKs are components of the mitogen activated protein kinase (MAPK) pathway and are believed to regulate small GTP-binding proteins (CDC42 and RAC) (Bagrodia and Cerione, 1999; Frisch, 2000) (see Fig. 12).

By probing scRNA-seq libraries we found that *PAK1* was widely expressed at relatively high levels in resting MG, and levels were significantly reduced in MG at different times after NMDA, and further reduced in MGPCs (Fig. 6a,b). Similarly, levels of *PPP2CA*, *CDC42, GIT1* and *ILK* were expressed at relatively high levels in resting MG, but were widely expressed at reduced levels in MGPCs and activated MG in damaged retinas (Fig. 6a,b). In addition, *PPP2CA, CDC42, GIT1* and *ILK* were expressed by different types of retinal neurons, NIRG cells and oligodendrocytes (Fig. 6a). We found that MG expressed *ITGB1* and other integrin isoforms, including *ITGA1, ITGA2, ITGA3* and *ITGA6* (Fig. 6a,b). In general, integrins were expressed at high levels in scattered resting MG, whereas levels were reduced, but more widely expressed among activated MG, and further reduced in MGPCs (Fig. 6a,b).

**Figure 6.**
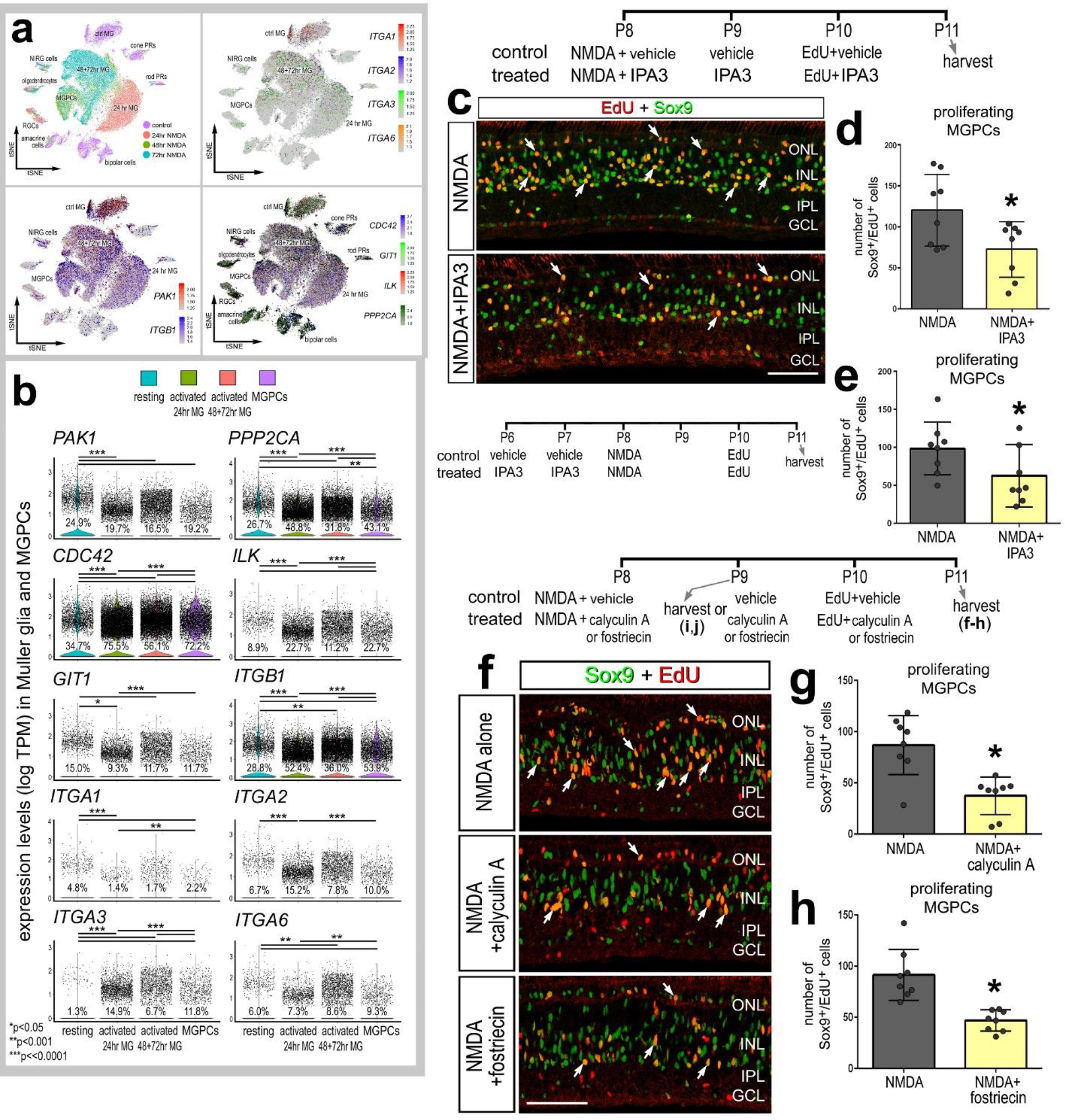
Patterns of expression and inhibition of putative MDK receptors, integrins, and signal transducers in damaged retinas. scRNA-seq libraries (Fig. 2) were probed for patterns of expression of integrin alpha/beta isoforms and associated signaling ITGB1 molecules p21 associated kinase-1 (PAK1), protein phosphatase 2a catalytic subunit alpha (PPP2CA), integrin linked kinase (ILK), and ARF GTPase-activating protein (GIT1). tSNE plots demonstrate patterns of expression of *PAK1, ITGB1, ITGA1, ITGA2, ITGA3, ITGA6, ITGAV, CAT, CDC42, GIT1, ILK* and *PPP2CA* (**a**). Violin/scatter plots indicate significant differences (*p<0.01, **p<0.001, ***p<<0.001; Wilcox rank sum with Bonferoni correction) in expression of *PAK1, ITGB1, ITGA1, ITGA2, ITGA3, ITGA6, ITGAV, CAT, CDC42, GIT1, ILK* and *PPP2CA* among MG and MGPCs (**b**). The number on each violin indicates the percentage of expressing cells. PAK1-specific inhibitor IPA3 was injected with and following NMDA (**c**,**d**) or before NMDA (**e**) and analyzed for proliferation of MGPCs. Alternatively, PP2A-specific inhibitors calyculin A or fostriecin were injected with and following NMDA (**f**-**h**). Sections of the retina were labeled for EdU (red) and Sox9 (green; **c, f**). Arrows indicate nuclei of proliferating MGPCs (**a,g**). The histogram/scatter-plots in **d, e, g** and **h** illustrate the mean (±SD) number of labeled cells. Each dot represents one biological replicate. Significance of difference (*p<0.05) was determined by using a paired *t*-test. Arrows indicate nuclei of proliferating MGPCs (**c,f**). The calibration bar panels **c** and **f** represent 50 μm. Abbreviations: ONL – outer nuclear layer, INL – inner nuclear layer, IPL – inner plexiform layer, GCL – ganglion cell layer, ns – not significant.

We next tested how PAK1 and PP2A influence the formation of MGPCs in NMDA-damaged retinas. MDK-signaling is known to be modulated by the second messenger PAK1 which is up-regulated in proliferating cancerous cells (Kumar et al., 2006). IPA3 is an isoform-specific allosteric inhibitor of PAK1 which prevents auto-phosphorylation (Deacon et al., 2008). Administration of IPA3 with NMDA significantly decreased numbers of proliferating MGPCs (Fig. 6c,d). By contrast, IPA3 had no significant effect upon the proliferation and accumulation of NIRG cells or microglia (Supplemental Fig. 4). Unlike Na_3_VO_4_, IPA3 had no impact on numbers of TUNEL^+^ cells compared to those seen in retinas treated with NMDA alone (Supplemental Fig. 4). Since *PAK1* expression was most prevalent in resting MG and decreased after NMDA damage, we tested whether application of IPA3 prior to NMDA influenced glial cells and neuronal survival. We found that IPA3 prior to NMDA resulted in a significant decrease in proliferating MGPCs (Fig. 6e), whereas there was no significant difference in numbers of dying cells or proliferation of microglia and NIRG cells (Supplemental Fig. 4). Similar to the effects of IPA3, two different inhibitors to PP2A, fostriecin and calyculin A, significantly decreased numbers of proliferating MGPCs in NMDA-damaged retinas (Fig. 6f-h). Fostriecin and calyculin A had relatively little effect upon the accumulation, reactivity, cell death and proliferation of NIRG cells and microglia, with the exception of a small but significant decrease in proliferating microglia with calyculin A-treatment compared to controls (Supplemental Fig. 4). Collectively, these findings suggest that inhibition of signal-transducers of MDK-signaling suppresses the formation of MGPCs.

### *MDK* in retinas treated with insulin and FGF2

In the postnatal chick retina, the formation of proliferating MGPCs can be induced by consecutive daily injections of Fibroblast growth factor 2 (FGF2) and insulin in the absence of neuronal damage (Fischer et al., 2002b). Eyes were treated with two or three consecutive daily doses of FGF2 and insulin and retinas were processed to generate scRNA-seq libraries. Cells were clustered based on their gene expression in UMAP plots and colored by their library of origin (Fig. 7a,b). MG glia were identified based on collective expression of *VIM, GLUL* and *SLC1A3* and MGPCs were identified based on expression of *TOP2A, NESTIN, CCNB2* and *CDK1* for MGPCs (Supplemental Fig. 5a,b). Resting MG from saline-treated retinas formed a cluster distinct from MG from retinas treated with two- and three-doses of FGF2+insulin based on unique patterns of gene expression (Fig. 7b; Supplemental Fig. 5a,b). Additionally, MG treated with 2 versus 3 doses of insulin and FGF2 were sufficiently dissimilar to follow different trajectories of gene expression in pseudotime analysis (Supplemental Fig. 5c,d).

**Figure 7.**
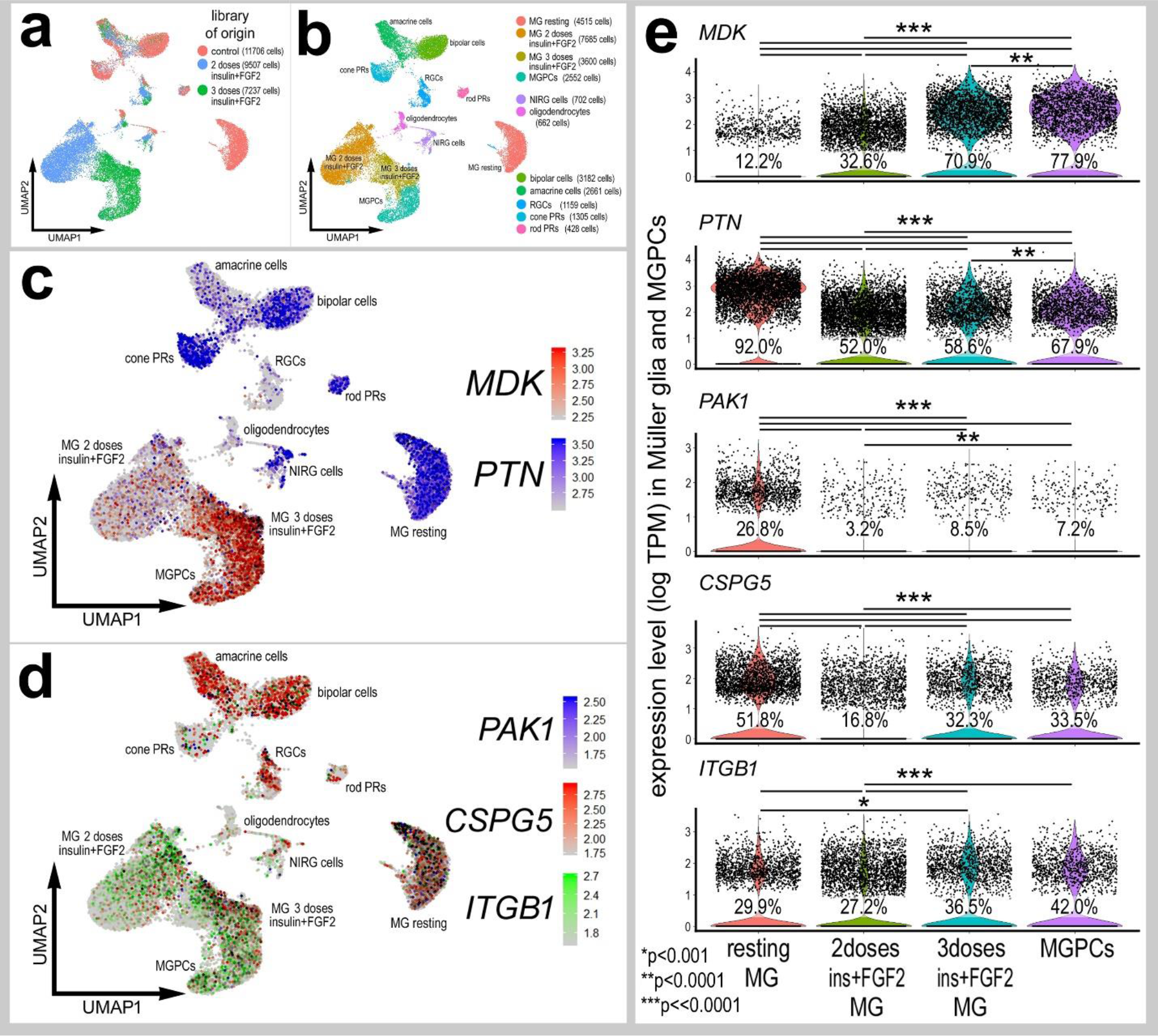
MG expression of *MDK* and putative MDK-receptors in retinas treated with insulin and FGF2. scRNA-seq was used to identify patterns of expression of MDK-related genes among cells in saline-treated retinas and in retinas after 2 and 3 consecutive doses of FGF2 and insulin (**a,b**). In UMAP plots, each dot represents one cell, and expressing cells indicated by colored heatmaps of gene expression for *MDK, PTN, PAK1, CSPG5* and *ITGB1* (**c,d**). Black dots indicate cells with expression of two or more genes. (**e**) Changes in gene expression among UMAP clusters of MG and MGPCs are illustrated with violin plots and significance of difference (*p<0.1, **p<0.0001, ***p<<0.0001) determined using a Wilcox rank sum with Bonferoni correction. The number on each violin indicates the percentage of expressing cells.

Similar to patterns of express in NMDA-damaged retinas, there was a significant increase in *MDK* with growth factor-treatment as demonstrated by patterns of expression in UMAP and violin plots, and pseudotime analyses (Fig. 7c-e; Supplemental Fig. 5d-f). By comparison, levels of *PTN* were significantly decreased in MG following treatment with insulin and FGF2 (Fig. 7c,e; Supplemental Fig. 5d-f). Similarly, levels of *PAK1* were decreased in activated MG and MGPCs in response to growth factor treatment (Fig. 7d,e; Supplemental Fig. 5e,f). *CSPG5* was widely expressed at high levels in scattered resting MG, and was significantly reduced in MG and MGPCs following treatment with insulin and FGF2 (Fig. 7d,e; Supplemental Fig. 5e, f). *ITGB1* expression was scattered in resting MG, and decreased slightly in MG and MGPCs treated with insulin and FGF2 (Fig. 7d,e; Supplemental Fig. 5e,f). In sum, treatment with FGF2 and insulin in the absence of retina damage influenced patterns of expression for MDK-related genes similar to those seen in NMDA-damaged retinas.

We next isolated MG, aggregated and normalized scRNA-seq data from saline-, NMDA-, FGF2+insulin- and NMDA/FGF2+insulin-treated retinas to directly compare levels of *MDK, PTN* and related factors. UMAP plots revealed distinct clustering of MG from control retinas and MG from 24hrs after NMDA-treatment, whereas MG from retinas at 48 and 72hrs after NMDA and from retinas treated with insulin and FGF2 formed a large cluster with distinct regions (Fig. 8a-e). UMAP and Dot plots revealed distinct patterns of expression of genes associated with resting MG, de-differentiating MG, activated MG and proliferating MGPCs (Fig. 8c-e). Different zones, representing MGPCs in different phases of the cell cycle were comprised of cells from different times after NMDA-treatment and FGF2+insulin-treatment (Fig. 8e). Expression of *MDK* was most widespread and significantly up-regulated in MG in damaged retinas and MGPCs compared to MG from retinas treated with insulin and FGF2 (Fig. 8f). Compared to levels seen in resting MG, levels of *PTN* were reduced in activated MG from damaged retinas and MGPCs, and levels were further decreased in MG from normal and damaged retinas that were treated with insulin and FGF2 (Fig. 8f); similar patterns of expression were seen for *PAK1, CSPG5, ITGB1, PPP2CA* and *CDC42.* By contrast, levels of *SDC4* were highest and most widespread in MG at 24hrs after NMDA-treatment and were relatively reduced in all other groups of MG and MGPCs (Fig. 8f). Collectively, these findings indicate that damaged-induced changes of *MDK, PTN* and related factors in MG are very dramatic, and these changes in relative expression levels in MG are dampened by insulin and FGF2 whether applied to undamaged or damaged retinas.

**Figure 8.**
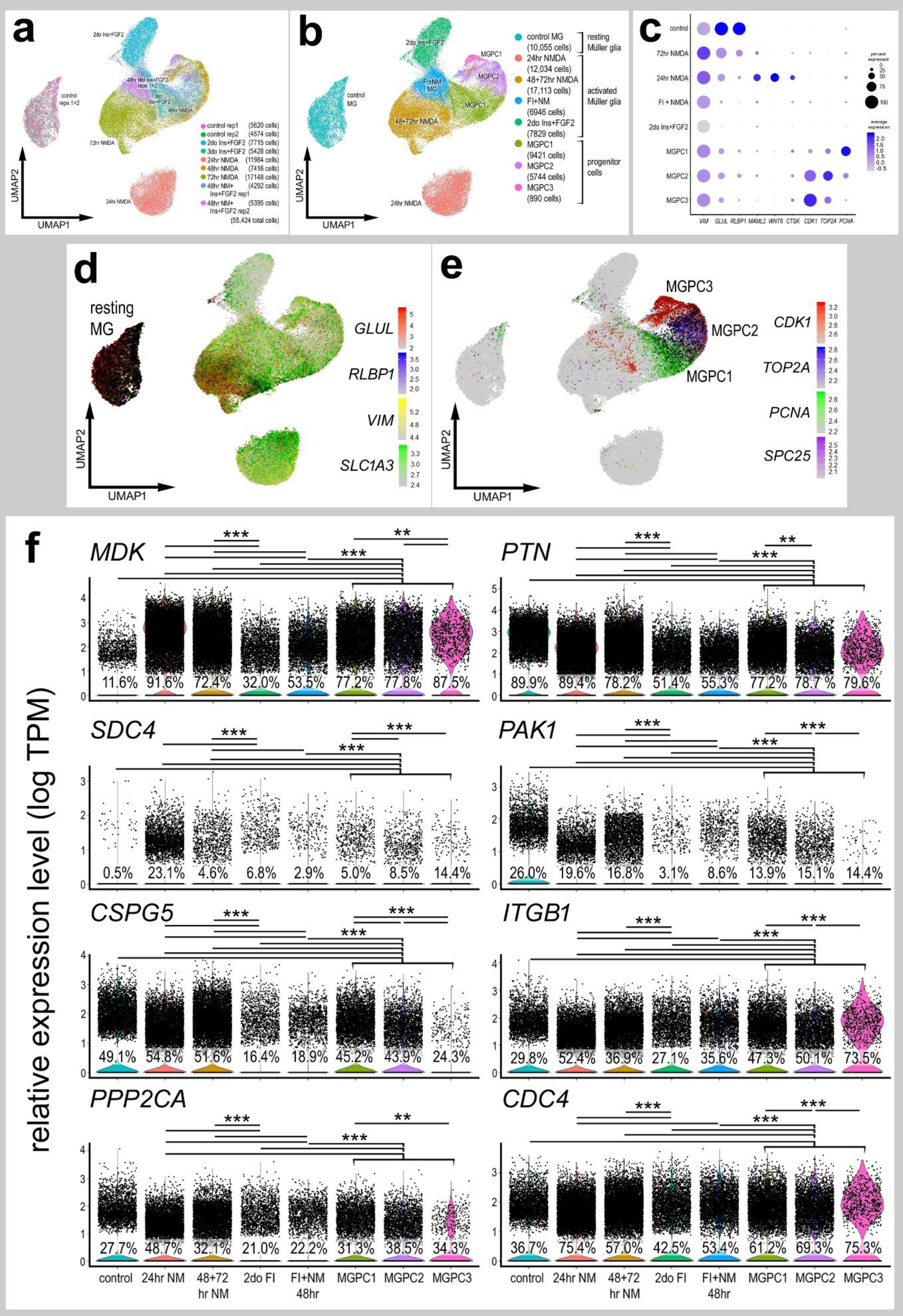
Expression of *MDK* and MDK-related genes in aggregate of scRNA-seq libraries for MG from different treatments. In UMAP and violin plots each dot represents one cell. MG were bioinformatically isolated from 2 biological replicates for control retinas and retinas treated with 2 doses of insulin and FGF2, 3 doses of insulin and FGF2, 24 hrs after NMDA, 48 hrs after NMDA, 48hrs after NMDA + insulin and FGF2, and 72 hrs after NMDA. UMAP analysis revealed distinct clusters of MG which includes control/resting MG, activated MG from retinas 24hrs after NMDA treatment, activated MG from 2 doses of insulin and FGF2, activated MG from 3 doses of insulin FGF2 and NMDA at different times after treatment, activated MG returning toward a resting phenotype from 48 and 72 hrs after NMDA-treatment, and 3 regions of MGPCs. The dot plot in **c** illustrates some of the pattern-distinguishing genes and relative levels across the different UMAP-clustered MG and MGPCs. UMAP plots illustrate the distinct and elevated expression of *GLUL, RLBP, VIM* and *SLC1A3* in resting MG (**d**) and *CDK1, TOP2A, PCNA* and *SPC25* in different regions of MGPCs (**e**). Violin plots in **f** illustrate relative expression levels for *MDK, PTN, SDC4, PAK1, CSPG5, ITGB1, PPP2CA* and *CDC4* in UMAP-clustered MG and MGPCs. Significance of difference (**p<0.001, ***p<<0.001) was determined by using a Wilcox rank sum with Bonferoni correction. The number on each violin indicates the percentage of expressing cells.

### MDK and MGPCs in undamaged retinas

In the absence of damage, in retinas treated with FGF2 and insulin, we tested whether MDK influences MG and the formation of MGPCs. MDK was administered two days before and with three consecutive daily doses of FGF2 and insulin (Fig. 9). MDK-treatment had no significant influence upon the formation of proliferating MGPCs or the accumulation of NIRG cells (Fig. 9a,b,c). We next tested whether inhibition of PAK1 with IPA3 influenced glial cells in retinas treated with FGF2 and insulin. IPA3-treatment had no significant influence upon the formation of proliferating MGPCs, NIRG cells or microglia (Fig. 9d-g), but did have a small, but significant, inhibitory effect upon the accumulation of NIRG cells (Fig. 9h). We next tested whether inhibition of PTPRZ with Na_3_VO_4_ influenced the glial cells in retinas treated with FGF2 and insulin. There was no significant difference in numbers of proliferating MGPCs following 3 days of treatment with insulin and FGF2 with Na_3_VO_4_ (Fig. 9i). Similarly, MDKi inhibitor had no effect upon the formation of proliferating MGPCs (not shown). However, there was an increase in the total number of NIRG cells in the retina in response to Na_3_VO_4_ treatment (Fig. 9j-k). The reactivity and accumulation of microglia were unaffected by the Na_3_VO_4_ or IPA3 (data not shown).

**Figure 9.**
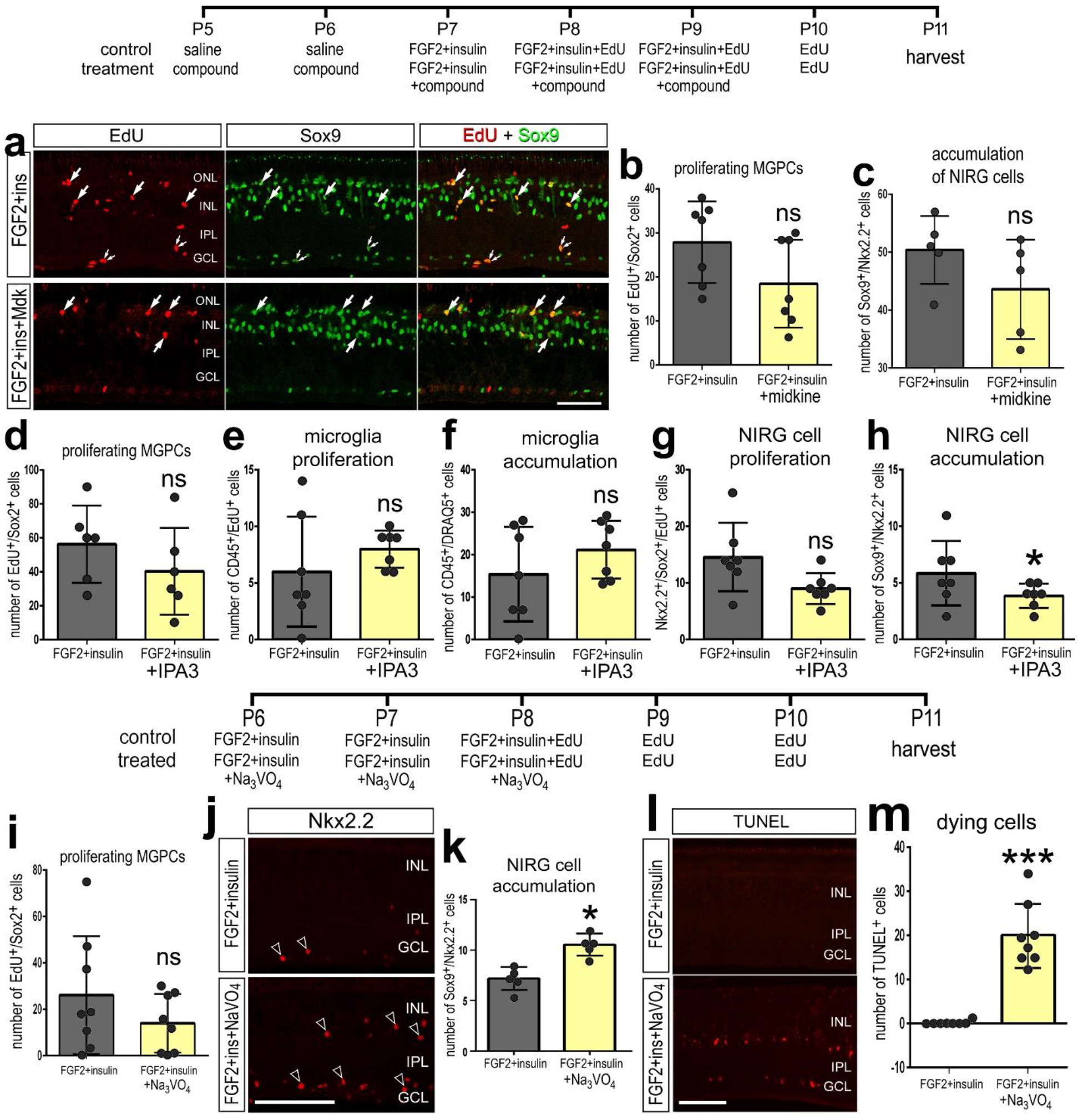
The combination of MDK with FGF2 and insulin had no effect on the formation of proliferating MGPCs. (**a**-**h**) Retinas were obtained from eyes that received 2 consecutive daily doses of vehicle or MDK/IPA3, 3 consecutive daily injections of FGF2 and insulin with or without MDK/IPA3, followed by an injection of EdU, and harvested 24 hrs after the final injection. (**i**-**j**) Alternatively, retinas were obtained from eyes that received, 3 consecutive daily injections of FGF2 and insulin with or without NaVO4, followed by 2 consecutive daily injections of EdU, and harvested 24 hrs after the final injection. The responses of MG, microglia, and NIRG proliferation and accumulation was evaluated after harvesting. Sections of the retina were labeled for EdU (red; **a**), TUNEL(**l**) or antibodies to Sox9 (green; **a**, **j**) and Nkx2.2 (red; **j**). The histogram/scatter-plots illustrate the mean (±SD) number of proliferating MGPCs, microglia, NIRG cells, or TUNEL positive nuclei. Significance of difference (*p<0.05) was determined by using a *t*-test. Arrows indicate EdU^+^/Sox9^+^ MG, small double-arrows indicate proliferating NIRG cells, and hollow arrow-heads indicate Sox9^+^/Nkx2.2^+^ NIRG cells. The calibration bars in panels **a** and **j** represent 50 μm. Abbreviations: ONL – outer nuclear layer, INL – inner nuclear layer, IPL – inner plexiform layer, GCL – ganglion cell layer.

### *Mdk, Ptn* and MDK-receptors in damaged mouse retinas

We next sought to assess the expression of *Mdk* and related factors in normal and NMDA-damaged mouse retinas. Comparison of the different responses of glial cells across species can indicate important factors that confer the potential of MG to reprogram into MGPCs (Hoang et al., 2019). UMAP analysis of cells from control and NMDA-damage mouse retinas revealed discrete clusters of different cell types (Fig. 10a). Neuronal cells from control and damaged retinas were clustered together, regardless of time after NMDA-treatment (Fig. 10a). By contrast, resting MG, which included MG from 48 and 72 hrs after NMDA, and activated MG from 3, 6, 12 and 24 hours after treated were spatially separated in UMAP plots (Fig. 10a,b). Pseudotime analysis placed resting MG (control and some MG from 48 and 72 hrs after treatment) to the left, MG from 3 and 6 hrs after treatment to the far right, and MG from 12 and 24 hrs bridging the middle (Supplemental Fig. 6a-d). Unlike chick MG, mouse MG rapidly downregulate *Mdk* in response to damage and this downregulation is maintained through 72 hrs after treatment (Fig. 10c,d). Similar to MG in the chick, *Ptn* was rapidly and significantly down-regulated at 3hrs, and further down-regulated at 6hrs, and sparsely expressed at 12-48hrs (Fig. 10c,d). Levels of *Pak1* were low in resting MG, and elevated in MG only at 3hrs after NMDA-treatment (Fig. 10c,d). Similar to chick MG, *Cspg5* was significantly decreased in activated MG in damaged retinas (Fig. 10c,d). By contrast, there were significant increases in levels of *Sdc4* and *Itgb1* in MG in damaged retinas (Fig. 10c,d; Supplemental Fig. 6e-g). We further analyzed the responses of MG in damaged retinas at 48hrs after NMDA ± treatment with insulin and FGF2, which is known to stimulate proliferation of MG (Karl et al., 2008). Treatment with FGF2 and insulin in damaged retinas significantly reduced levels of *Glul,* whereas levels of *Vim* and *Gfap* were significantly increased (Supplemental Fig. 6h-j). By comparison, levels of *Mdk* and *Sdc4* were significantly increased in MG in retinas treated with NMDA+FGF2/insulin, whereas levels of *Ptn*, *Cspg5* and *Itgb1* were unchanged (Supplemental Fig. 6h-j).

**Figure 10.**
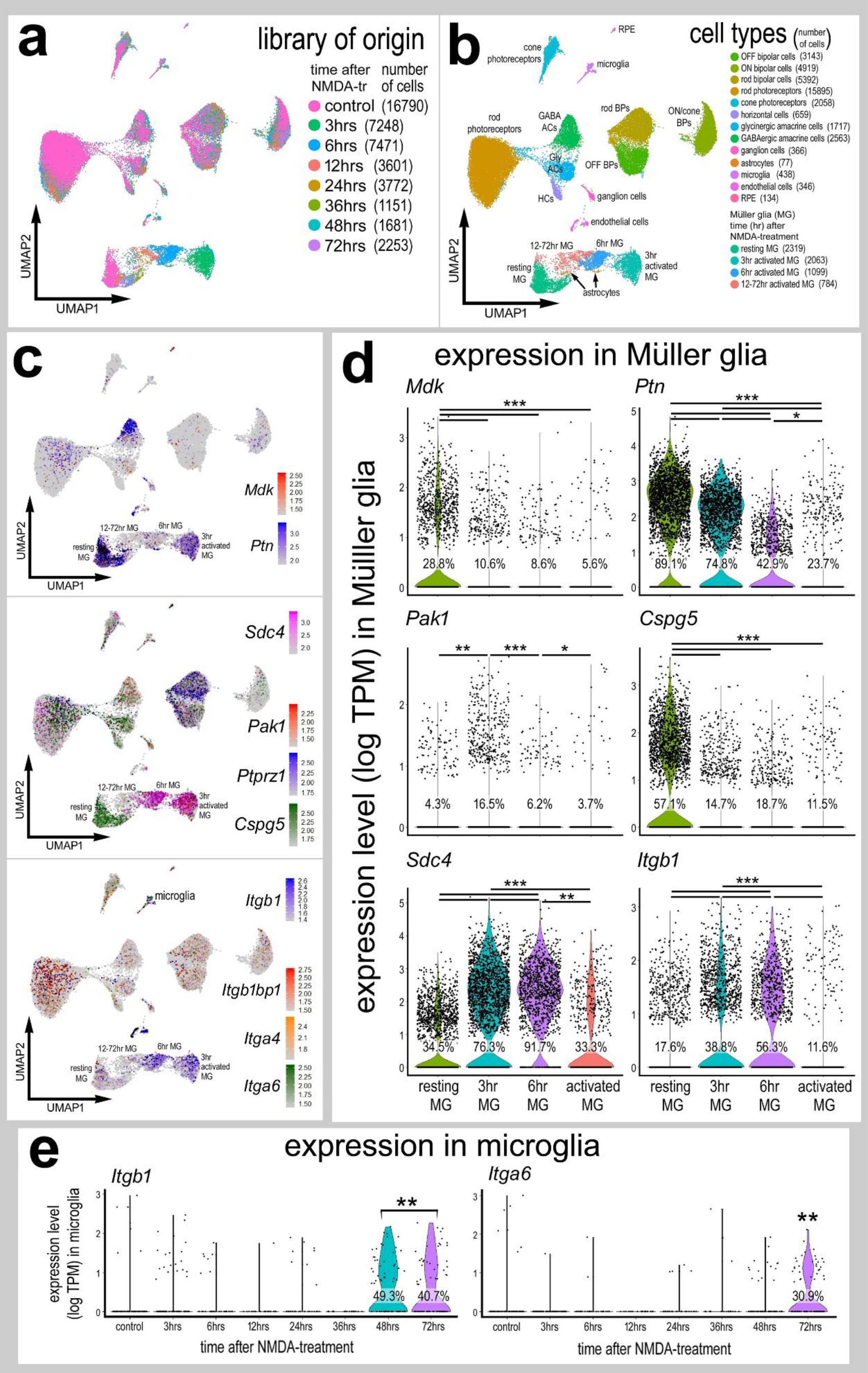
Mouse MG have a reduced profile of *Mdk, Ptn* and MDK-related genes in MG in response to NMDA damage. Cells were obtained from control retinas and from retinas at 3, 6, 12, 24, 36, 48 and 72hrs after NMDA-treatment and clustered in UMAP plots with each dot representing an individual cell (**a**). UMAP plots revealed distinct clustering of different types of retinal cells; resting MG (a mix of control, 48hr and 72hr NMDA-tr), 12-72 hr NMDA-tr MG (activated MG in violin plots), 6hrs NMDA-tr MG, 3hrs NMDA-tr MG, microglia, astrocytes, RPE cells, endothelial cells, retinal ganglion cells, horizontal cells (HCs), amacrine cells (ACs), bipolar cells (BPs), rod photoreceptors, and cone photoreceptors (**b**). Cells were colored with a heatmap of expression of *Mdk, Ptn, Sdc4, Pak1, Ptprz1, Cspg5, Itgb1bp1, Itga4* and *Itgba6* gene expression (**c**). Black dots indicates cells with two or more markers. In MG, changes in gene expression are illustrated with violin/scatter plots of *Mdk, Ptn, Pak1, Cspg5, Sdc4, and Itgb1* and quantified for significant changes (**d**) (*p<0.01, **p<0.0001, ***p<<0.001). Similarly, UMAP-clustered microglia were analyzed and genes *Itgb1* and *Itga6* were detected and quantified in violin plots for cells from each library of origin (**e**). The number on each violin indicates the percentage of expressing cells.

We next investigated the activation of different cell-signaling pathways in retinal cells in response to intravitreal delivery of MDK. We failed to detect activation of NFkB, pStat3, pSmad1/5/8, pCREB, p38 MAPK or pERK1/2 (not shown). However, in response to a single injection of MDK, we found a selective and significant up-regulation of cFOS and pS6 in MG (Fig. 11a-e), similar to that observed in chick retina. Other types of retinal cells did not appear to respond to MDK with up-regulation of cFOS or pS6. We next tested whether intraocular injections of MDK combined with NMDA-induced damage influences the proliferation of MG in the mouse retina. Consistent with previous reports (Karl et al., 2008), there were very few proliferating MG in NMDA-damaged retinas (Fig. 11f,h). By contrast, application of MDK with NMDA resulted in a small, but significant increase in numbers of proliferating MG (Fig. 11f-h).

**Figure 11.**
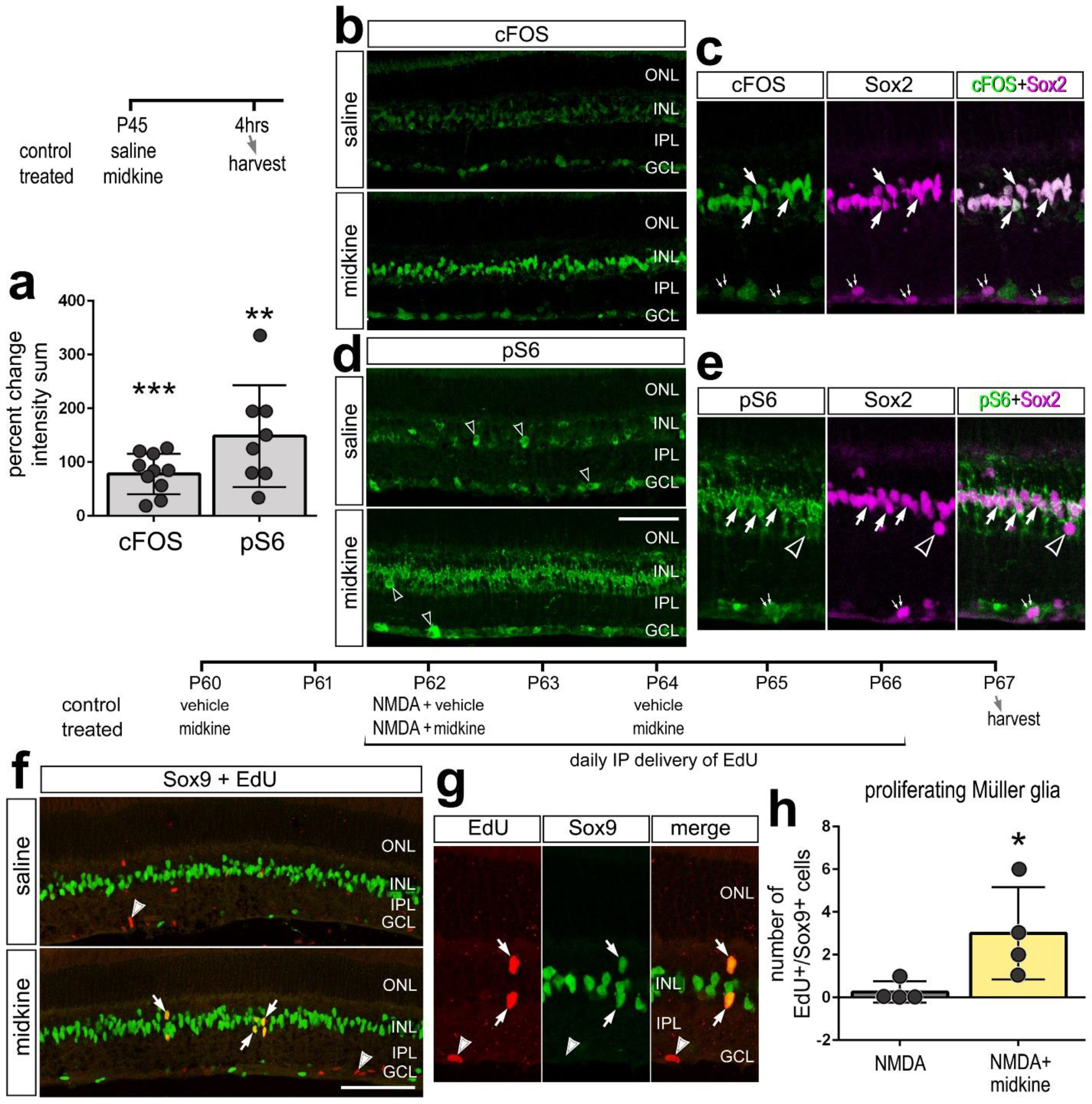
MDK activates cell-signaling in MG and stimulates proliferation in the mouse retina. (**a**-**e**) A single intraocular injection of MDK was delivered and retinas were harvested 4 hours later. The histogram in **a** represents the mean percent change (±SD) in density sum and area for percentage change in intensity sum for cFOS and pS6 immunofluorescence. Each dot represents one replicate retina. Significance of difference (**p<0.01, ***p<0.0001) was determined by using a paired two-way *t*-test. Vertical sections of saline (control) and MDK-treated retinas were labeled with antibodies to cFOS (green; **b**,**c**), pS6 (green; **d**,**e**) and Sox2 (magenta; **c**,**e**). (**f**-**h**) Treatment included intraocular injections of MDK or vehicle at P60, NMDA and MDK/vehicle at P62, MDK or vehicle at P60, Edu was applied daily by intraperitoneal (IP) injections from P62 through P66, and tissues were harvested at P67. The histogram in **h** represents the mean (±SD) numbers of EdU^+^/Sox9^+^ cells in the INL. Each dot represents one replicate retina. Significance of difference (*p<0.05) was determined by using a paired two-way *t*-test. Arrows indicate the nuclei of MG and arrow-heads indicate EdU^+^/Sox9^-^ cells (presumptive proliferating microglia). The calibration bar (50 μm) in panel **d** applies to **b** and **d**. Abbreviations: ONL – outer nuclear layer, INL – inner nuclear layer, IPL – inner plexiform layer, GCL – ganglion cell layer.

**Figure 12.**
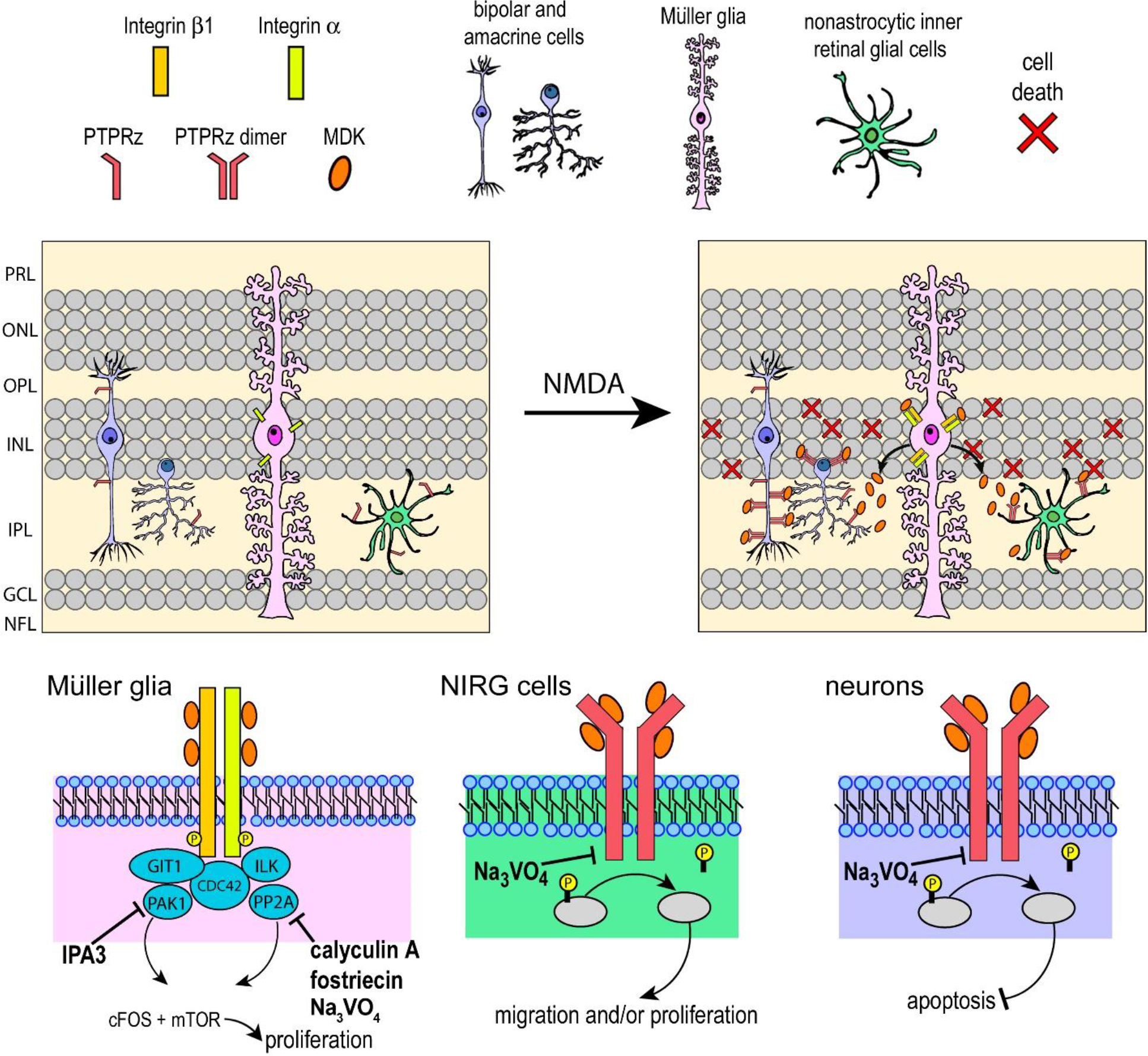
Schematic summary of MDK-signaling in normal and NMDA-damaged retinas. Patterns of expression, determined by scRNA-seq, are shown for Integrin β1, Integrin α, PTPRZ1, PAK1 and MDK in MG, NIRG cells and inner retinal neurons. Although GIT1, ILK, CDC42, and PP2A (*PPP2CA*) were widely expressed by nearly all retinal cells (according to scRNA-seq data; see Fig 6a), signaling through Integrins is shown only in MG because *ITG’s* were largely confined to MG. Putative sites of action are shown for small-molecule inhibitors, including IPA3, calyculin A, fostriecin and Na_3_VO_4_. Abbreviations: PRL – photoreceptor layer, ONL – outer nuclear layer, INL – inner nuclear layer, IPL – inner plexiform layer, GCL – ganglion cell layer, NFL – nerve fiber layer.

## Discussion

In the chick retina, we find that dynamic expression of *MDK* during retinal development and in mature retinas following injury or growth factor-treatment. High levels of *MDK* expression were selectively and rapidly induced in MG following damage or treatment with insulin and FGF2, with larger increases in expression seen in damaged tissues. Addition of exogenous MDK before damage was neuroprotective and resulted in decreased numbers of proliferating MGPCs. Antagonism of MDK-signaling reduced numbers of proliferating MGPCs and stimulated the accumulation of NIRG cells and increased numbers of dying cells. Na_3_VO_4_ and PAK1 antagonist had differential effects on NIRG cells and cell death that were context dependent. In contrast to the findings in chick, we find that *Mdk* is down-regulated by MG in damaged mouse retinas. In both chick and mouse retinas, exogenous MDK selectively induces mTOR-signaling and expression of cFOS in MG.

### PTN signaling in the retina

PTN and MDK are in the same family of growth factors and are both dynamically expressed in the developing, damaged, and growth factor treated retinas. Although treatment with MDK had varying effects on MG, microglia, NIRGs, and neurons, PTN administration failed to illicit detectable effects upon retinal cells. Levels of *PTN* are high in resting MG and down-regulated in response to neuronal damage or treatment with insulin and FGF2. In principle, PTN acts at the same receptors as MDK, but expression of receptor isoforms may underlie the different cellular responses to MDK and PTN. PTN has preferred binding-affinity for SDC4 and ITGB3, whereas MDK has preferred binding-affinity for SDC3 and ITGB1 (Muramatsu et al., 2004; Raulo et al., 1994; Xu et al., 2014), which are expression by bipolar cells and MG.

PTN and MDK may induce different biological effects on the same receptors. For instance, data suggests that there may be differential receptor activity between PTN and MDK on the PTPRz receptor. Binding of PTN to PTPRz induces oligomerization of the receptor that reduces phosphatase activity (Fukada et al., 2006). Conversely, MDK promotes embryonic neuronal survival in a PTPRz receptor complex, which is inhibited by Na_3_VO_4_ (Sakaguchi et al., 2003). Although we failed to detect PTN-mediated effects upon retinal cells, PTN may serve other important biological roles in retinal hometostasis, glial phenotype/functions or neuroprotection in other models of retinal damage.

### Receptor expression and cells responding to MDK

The effects of MDK on retinal glia has not been studied in mammals or birds. In acutely damaged chick retina, MG are capable of forming numerous proliferating progenitor cells (MGPCs), but few of the progeny differentiate into neurons (Fischer and Reh, 2001; Fischer and Reh, 2002). The reprogramming of MG into MGPCs can be induced by FGF2 and insulin in the absence of damage through MAPK signaling (Fischer and Reh, 2002; Fischer et al., 2002a; Fischer et al., 2002b). Similarly, IGF1, BMP, retinoic acid, sonic hedgehog, Wnt, and Jak/Stat agonists have been observed to enhance the formation of MGPCs (Fischer et al., 2009a; Gallina et al., 2015; Todd and Fischer, 2015; Todd et al., 2016; Todd et al., 2017; Todd et al., 2018). Consistent across the different signaling pathways that drive the formation of proliferating MGPCs is the up-regulation of cFOS and necessity for mTor-signaling in reprogramming MG (Zelinka et al., 2016). Previous reports have provided many examples of MDK activating cell-signaling pathways that drive proliferation (Reiff et al., 2011; Winkler and Yao, 2014). Accordingly, we propose that MDK-mediated cell-signaling that results in activation of cFOS and mTOR contributes to the network of pathways that drive the formation of proliferating MGPCs in the chick retinas.

Patterns of expression for receptors suggests than glial cells and inner retinal neurons are targets of MDK and PTN. In MG the predominant receptor is *ITGB1,* whereas NIRGs cells express *PTPRZ1,* and amacrine and bipolar cells express a combination of *PTPRZ1* and *SDC4.* The mechanism by which ITGB1 and PTPRZ influence cell cycle and differentiation are distinctly different. PTPRZ promotes stem cell characteristics and ligand binding inhibits this phosphatase function (Fujikawa et al., 2016; Fukada et al., 2006; Kuboyama et al., 2015). Signal transduction through ITGB1 influences cytoskeleton remodeling that is associated with cell migration and proliferation (Muramatsu et al., 2004). ITGB1 can activate or inhibit secondary messengers depending on tyrosine phosphorylation (Kim et al., 2004; Mulrooney et al., 2000; Song et al., 2014). Ligand binding to ITGB1 initiates tyrosine phosphorylation of intracellular domains, and integrin linked kinases (ILKs) activate PP2A and cell cycle kinases, such as CDC42 (Ivaska et al., 1999; Ivaska et al., 2002). The transcriptional profiles of individual retinal cell types suggest that MDK and PTN likely have autocrine and paracrine actions that are dynamically regulated in damaged retinas and manifested through MG, and dynamic regulation of mRNA is strongly correlated with changes in protein levels and function (Liu et al., 2016).

### MDK-signaling in MG

Application of MDK prior to NMDA-induced damage decreased numbers of proliferating MGPCs and decreased numbers of dying cells. Levels of retinal damage positively correlate to the proliferative response of MG (Fischer and Reh, 2001; Fischer et al., 2004). We propose that the neuroprotective effects of MDK secondarily influenced the proliferative of MGPCs. It is possible that the addition of MDK to damaged retinas failed to influence MGPCs because of “ceiling effects” wherein (i) ligand/receptor interactions are saturated, (ii) the activity of secondary messengers are saturated, or (iii) the massive up-regulation of MDK by MG is not directly involved in driving the formation of proliferating MGPCs.

Phosphatase inhibitor Na_3_VO_4_ suppressed the formation of MGPCs and increased cell death, and these effects where blocked by addition of MDK. Inhibition of intracellular phosphatases, such as PP2A that are commonly associated with ITGB1 receptors, increasing the upstream activation may be attributed to restoring MG responses to damage. Inhibition of Git/Cat-1/PAK1-signaling is associated with ITGB1-mediated cytoskeleton remodeling during migration and proliferation (Martin et al., 2016; Muramatsu et al., 2004). Consistent with these observations, we found that inhibition of PAK1 and PP2A effectively suppressed the formation of MGPCs in damaged retinas. Further studies are required to identify changes in phosphorylation and expression that are down-stream of PP2A activity.

MDK and cell-signaling inhibitors failed to have significant impacts upon MG and microglia in retinas treated with insulin and FGF2. Similar to NMDA-treatment, we see significant changes in expression levels of *MDK, PTN, PAK1* and *CSPG5,* suggesting that MDK-signaling is active in undamaged retinas treated with insulin and FGF2 (see Fig. 7). However, direct comparison of relative expression levels of *MDK* and related genes across all treatment groups indicated that: (i) although *MDK* is up-regulated with insulin and FGF2, levels are much less than those seen with NMDA alone, (ii) *SDC4* is modestly induced in MG by insulin and FGF2, and (iii) levels of *PAK1, CSPG5, ITGB1, PPP2CA* and *CDC4* are further down-regulated by insulin and FGF2 compared to levels seen with NMDA-treatment. The diminished levels of MDK-receptors and signal transducers in MG treated with insulin and FGF2, compared to levels in MG treated with NMDA, may underlie the absence of effects of exogenous MDK and inhibitors. This suggests that MDK and down-stream signaling are not required for the formation of MGPCs in retinas treated with insulin+FGF2. Alternatively, the cell-signaling pathways that are activated by MDK are the same as those activated by insulin+FGF2 and there is no net gain in second messenger activation in MG by combining these factors. This is unique because many signaling pathways that have been implicated in regulating the formation of MGPCs in the chick retina are active following NMDA-induced damage and treatment with insulin and FGF2. These pathways include MAPK (Fischer et al., 2009a; Fischer et al., 2009b), mTOR (Zelinka et al., 2016), Notch (Ghai et al., 2010; Hayes et al., 2007), Jak/Stat (Todd et al., 2016), Wnt/b-catenin (Gallina et al., 2015), glucocorticoid (Gallina, 2015), Hedgehog (Todd and Fischer, 2015), BMP/SMAD (Todd et al., 2017), retinoic acid (Todd et al., 2018) and NFkB-signaling (Palazzo et al., 2020).

### MDK signaling in NIRG cells

A well-established receptor of MDK is PTPRz (Maeda et al., 1999). PTPRz is a cell-surface receptor that acts as a protein tyrosine phosphatase and is known to promotes proliferation (Fujikawa et al., 2016). This receptor is activated by MDK (Sakaguchi et al., 2003), but is deactivated by the binding of PTN through dimerization and tyrosine phosphorylation (Kuboyama et al., 2015). NIRG cells predominantly express *PTPRZ1* and the accumulation of these cells in response to damage was decreased with MDK-treatment and increased by treatment with phosphatase inhibitor Na_3_VO_4_. The accumulation of NIRG cells may result, in part, from migration, as MDK has been associated with migration and process elongation in different cell types (Ichihara-Tanaka et al., 2006; Kuboyama et al., 2015; Qi et al., 2001). Given that nothing is currently known about the specific functions of NIRG cells, it is difficult to infer how activation/inhibition of MDK-signaling in these glia impacts the reprogramming of MG or function/survival of retinal neurons.

### MDK and reprogramming of MG into MGPCs

There has been significant research understanding the cell-signaling pathways involved in the reprogramming of MG into proliferating MGPCs. IGF1, BMP, retinoic acid, HB-EGF, sonic hedgehog, Wnt, and CNTF are known to enhance the formation of MGPCs (Fischer et al., 2009a; Fischer et al., 2009b; Gallina et al., 2015; Todd and Fischer, 2015; Todd et al., 2016; Todd et al., 2017; Todd et al., 2018). The roles of these different pathways are similar in chick and zebrafish models of retinal regeneration, despite different capacities for neurogenesis (Goldman, 2014; Wan and Goldman, 2016). MDK has been implicated as an important factor to drive de-differentiation MG into proliferating progenitor cells (Calinescu et al., 2009; Luo et al., 2012; Nagashima et al., 2019). In the chick model, different inhibitors to PAK1, PP2A and PTPRZ1 had relatively modest impacts on the formation of MGPCs. Although exogenous MDK likely added nothing to already saturated levels in damaged retinas, MDK alone was not sufficient to induce the formation of MGPCs in the absence of damage when levels of MDK were low. These findings suggest that MDK signaling is not a primary signaling component to drive the formation of MGPCs in chick, unlike the key role for MDK seen in zebrafish (Gramage et al., 2015; Nagashima et al., 2019). In the chick, our findings suggests that MDK has pleiotropic roles and serves to both minimize neuronal cell death after damage and regulate the accumulation of NIRG cells. By comparison to effects seen in chick MG, MDK stimulated mTor-signaling and cFos expression in mouse MG and induced a modest increase in numbers of proliferating MG in damaged retinas. The neurogenic potential of these few proliferating MG remains to be determined.

## Conclusions

*MDK* and *PTN* are highly expressed by maturing MG in embryonic retinas, and *MDK* is down-regulated while *PTN* remains highly expressed by resting MG in the retinas of hatched chicks. In mature mouse retinas, *PTN* had similar patterns of expression, whereas *MDK* showed patterns of expression opposite to those seen in chick MG. When MG are stimulated by growth factors or neuronal damage in the chick model, *MDK* is robustly up-regulated whereas *PTN* is down-regulated. Injections of MDK had significant effects upon proliferating glia, formation of MGPCs, and glial reactivity, whereas we failed to detect cellular responses to exogenous PTN. Elevated MDK demonstrated survival-promoting effects upon neurons, and subsequently suppressed MGPC formation. Inhibiting factors associated with the ITGB1 signaling complex dampened MGPC formation and phosphatase inhibitor Na_3_VO_4_ over-rode the effects of MDK upon neuronal survival and MGPC formation. This effect was limited to damaged retinas, whereas in retinas treated with insulin+FGF2 there may be a convergence or over-lap of cell-signaling pathways activated by MDK and insulin+FGF2. In determining its effectiveness at driving dedifferentiation and proliferation of MG in mouse models of excitotoxic damage, only a small but significant increase in MGPCs was observed. Overall, the up regulation of *MDK* is among the largest increases in gene expression detected in MG stimulate by damage or growth factors, implying significant multifactorial functions in the context of development, reprogramming, and response to tissue damage.

## Author contributions

WAC – experimental design, execution of experiments, collection of data, data analysis, construction of figures and writing the manuscript. MF and IP – execution of experiments and collection of data. TH and SB – preparation of scRNA-seq libraries. AJF – experimental design, data analysis, construction of figures and writing the manuscript.

## Data availability

RNA-Seq data are deposited in GEO (GSE135406) Chick scRNA-Seq data can be queried at https://proteinpaint.stjude.org/F/2019.retina.scRNA.html.

## Acknowledgements

This work was supported by RO1 EY022030-08 (AJF) and UO1 EY027267-04 (SB, AJF).

