## Supplemental Tables and Figures for "Midkine in chick and mouse retinas: neuroprotection, glial reactivity and the formation of Müller glia-derived progenitor cells"

**Table S1:** Compounds used in this study with sources and dosages. A dose is 2µl and 20µl in mice and chick retinal injections respectively.

| Drug | company | product number | dosage |
|---|---|---|---|
| NMDA | Sigma | M3262 | 1.9mM/dose |
| 5-ethynyl-2'-deoxyuridine (Edu) | ThermoFischer | A10044 | 2µg/dose |
| FGF2 | R&D Systems | 234-FSE | 250ng/dose |
| Insulin | Sigma | I6634 | 1µg/dose |
| Sodium Orthovanadate | Sigma | 450243 | 0.27mM/dose |
| IPA3 | Tocris Biosciences | 3622 | 0.7mM dose |
| Calyculin A | Sigma | C5552 | 0.5mM dose |
| Fostreicin | Sigma | F4425 | 1.1mM dose |
| recombinant chick Midkine | MyBioSource | MBS1040042 | 1ug/dose |
| recombinant mouse Midkine | sigma | SRP3301 | 1ug/dose |

**Table S2.** Antibodies, sources and working dilutions.

| antibody | company | product number | host | clonality | dilution |
|---|---|---|---|---|---|
| Sox9 | Millipore | AB5535 | rabbit | polyclonal | 1:1000 |
| SOX2 | R&D Systems | AF2018 | goat | polyclonal | 1:500 |
| cFos | Santa Cruz | K-25 | rabbit | polyclonal | 1:300 |
| pS6 (ser240/244) | Cell Signaling Technologies | 2215S | rabbit | polyclonal | 1:300 |
| CD45 | Cedi Diagnostics | HIS-C7 | mouse | polyclonal | 1:200 |
| pHisH3 | Sigma | H0412 | rabbit | polyclonal | 1:300 |
| AP2-alpha | Developmental Studies Hybridoma Bank | AB_2313948 | mouse | monocolonal (3B5) | 1:1000 |
| OTX2 | R&D Systems | AF1979 | goat | polyclonal | 1:500 |
| Nkx2.2 | Developmental Studies Hybridoma Bank | 74.5A5 | mouse | monocolonal (74.5A5) | 1:50 |

**Supplemental Figure 1**. Annotation of aggregate scRNA-seq of embryonic chick retinal cells with putative receptors. scRNA-seq was used to identify patterns of expression of MDK-related receptor genes among embryonic retinal cells. In UMAP plots (**a**-**c**), violin plots (**d**), pseudotime trajectories (**e**) and pseudotime plots (**f**), each dot represents one cell. Cells were obtained from retinas at different stages of development E5 (3716 cells), E8 (9310 cells), E12 (5320 cells), and E15 (4352 cells). UMAP plots revealed distinct clustering of different types of retinal cells; E5 retinal progenitor cells (RPCs; 5288 cells), late RPCs and immature Müller glia (iMG; 3354 cells), maturing Müller glia (mMG; 2665 cells), immature bipolar cells (iBPs; 3132 cells), maturing OFF bipolar cells (OFF BPs; 1430 cells), maturing ON bipolar cells (ON BPs; 1692 cells), retinal ganglion cells (RGCs; 701 cells), immature amacrine cells (iACs; 1038 cells), maturing amacrine cells (mAC; 1692 cells), rod photoreceptors (rod PR; 1210 cells), and cone photoreceptors (cone PR; 404 cells) (**a**). RPCs were identified based on collective expression of *ASCL1* and genes associated with proliferation such as *CDK1* and *TOP2A* (**b**). Mature MG were identified based on collective expression of *SLC1A3*, *GLUL* and *RLBP1* (**b**). Patterns of expression of *ITGB1, PAK1, PTPRZ1* and were determined in UMAP plots (**c**). The black dots demonstrate individual cells that exhibit expression of 2 or more markers. Violin/scatter plots indicate significant differences (*p<0.1, **p<0.0001, ***p<<0.0001; Wilcox rank sum with Bonferoni correction) in expression of *ITGB1, PAK1, PTPRZ1* and in RPCs, iMG and mMG (**d**). The number on each violin indicates the percentage of expressing cells. RPCs, iMG and mMG were re-embedded for unsupervised pseudotime ordering of cells which revealed trajectories with RPCs clustered to the left and maturing MG clustered to the right (**e**). RPCs were identified based on collective expression of *ASCL1, CDK1 and* associated genes. Mature MG were identified by *GLUL* and *RLBP1.* These states were used as endpoints for the creation of a pseudotime trajectory, with RPCs at relative time 0. Abbreviations: RPC – retinal progenitor cell, MG – Müller glia, iMG – immature Müller glia, mMG - mature Müller glia.

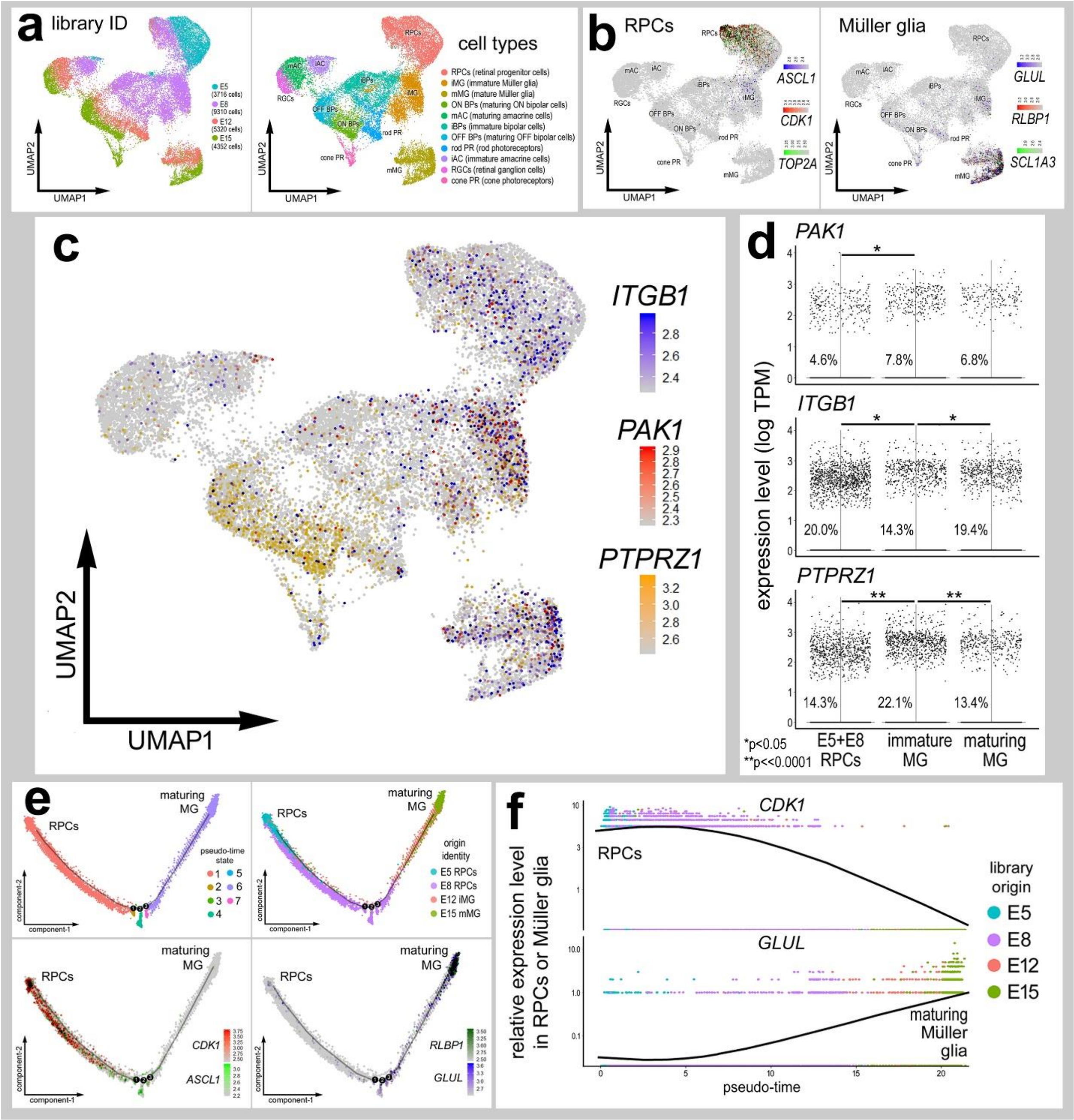
a
library ID
E5 (3716 cells)
E8 (9310 cells)
E12 (5320 cells)
E15 (4352 cells)
UMAP1
UMAP2
cell types
RPCs (retinal progenitor cells)
iMG (immature Müller glia)
mMG (mature Müller glia)
ON BPs (maturing ON bipolar cells)
mAC (maturing amacrine cells)
iBPs (immature bipolar cells)
OFF BPs (maturing OFF bipolar cells)
rod PR (rod photoreceptors)
iAC (immature amacrine cells)
RGCs (retinal ganglion cells)
cone PR (cone photoreceptors)
b
RPCs
ASCL1
CDK1
TOP2A
Müller glia
GLUL
RLBP1
SCL1A3
c
ITGB1
PAK1
PTPRZ1
UMAP1
UMAP2
d
PAK1
ITGB1
PTPRZ1
expression level (log TPM)
4.6%
7.8%
6.8%
20.0%
14.3%
19.4%
14.3%
22.1%
13.4%
*p<0.05
**p<<0.0001
E5+E8 RPCs
immature MG
maturing MG
e
RPCs
maturing MG
pseudo-time state
origin identity
E5 RPCs
E8 RPCs
E12 iMG
E15 mMG
CDK1
ASCL1
RLBP1
GLUL
component-1
component-2
f
CDK1
RPCs
GLUL
maturing Müller glia
relative expression level in RPCs or Müller glia
pseudo-time
library origin
E5
E8
E12
E15

**Supplemental Figure 2**. Expression of *MDK*, *PTN* and putative MDK-receptor genes in retinal cells following NMDA-treatment of the chick retina. scRNA-seq was used to identify patterns of expression of MDK-related genes among acutely dissociated retinal cells. In UMAP (**a**-**d**) and violin plots (**h**,**i**), each dot represents one cell. In UMAP plots black dots demonstrate individual cells that express 2 or more markers. Cells were obtained from control retinas (16,595 cells), and from retinas at 24hrs (14,918 cells), 48hrs (8483 cells) and 72 hrs (18,338 cells) after NMDA-treatment (**a**). UMAP plots revealed distinct clustering of different types of retinal cells; control MG (5468 cells), 24 hr NMDA-treated MG (12,576 cells), 48+72 hrs NMDA-treated MG (18,848 cells), MGPCs (5336 cells), microglia (66 cells), oligodendrocytes (1494 cells), NIRG cells (1224 cells), retinal ganglion cells (RGCs; 2049 cells), amacrine cells (2635 cells), bipolar cells (5169 cells), rod photoreceptors (974 cells), and cone photoreceptors (2495 cells) (**a**). MG were identified based on collective expression of *GLUL*, *RLBP1*, and *SLC1A3* and MGPCs were identified based on collective expression of *CDK1*, *ESPL1* and *TOP2A* (**b**). UMAP plots demonstrate patterns of expression of *CSPG5, PTN, PTPRZ1* (**c**), as includes presumptive MDK receptor and signaling genes *SDC4*, *PPP2CA*, and *ITGB1* (**d**). Violin plots illustrate expression of *PTN*, *CSPG5*, *PTPRZ1*, and *PPP2CA* among different cell populations after NMDA damage (**e**). Violin/scatter plots indicate significant differences (***p<<0.001; Wilcox rank sum with Bonferoni correction) in expression among MG and MGPCs (**f**). Violin/scatter plots indicate significant differences (**p<0.001, ***p<<0.0001; Wilcox rank sum with Bonferoni correction) in expression.

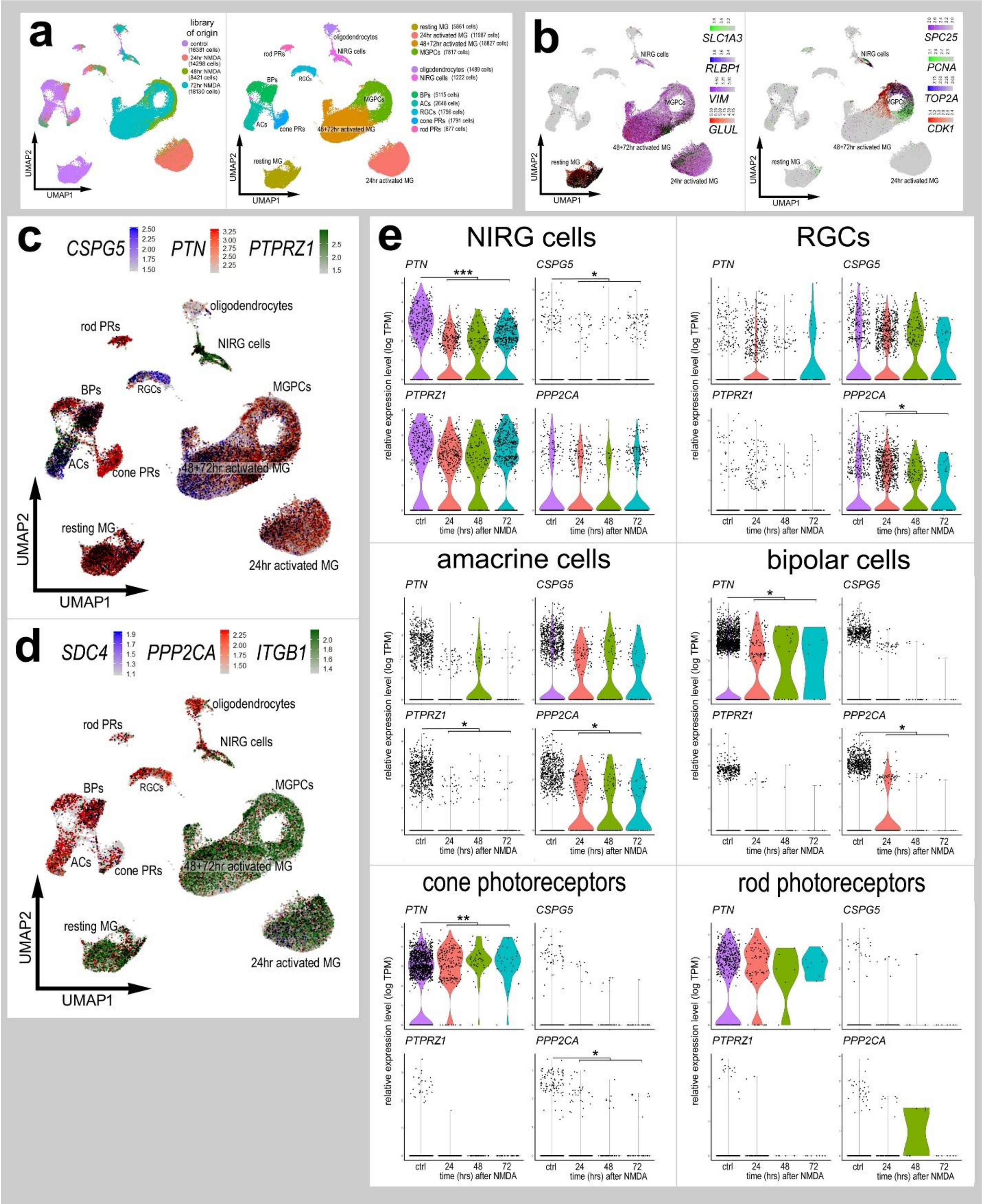
a
library of origin
control (16381 cells)
24hr NMDA (14298 cells)
48hr NMDA (8421 cells)
72hr NMDA (18130 cells)
UMAP1
UMAP2
resting MG
24hr activated MG
48+72hr activated MG
MGPCs
oligodendrocytes
NIRG cells
BPs
ACs
RGCs
cone PRs
rod PRs
b
SLC1A3
RLBP1
VIM
GLUL
SPC25
PCNA
TOP2A
CDK1
c
CSPG5
PTN
PTPRZ1
d
SDC4
PPP2CA
ITGB1
e
NIRG cells
RGCs
amacrine cells
bipolar cells
cone photoreceptors
rod photoreceptors
relative expression level (log TPM)
ctrl
24
48
72
time (hrs) after NMDA

**Supplemental Figure 3**. Pseudotime analysis and characterization of *MDK*, *PTN* and putative MDK-receptor genes in retinal cells following NMDA-treatment and NMDA plus FGF2 with insulin of the chick retina. The aggregate libraries were projected into an unbiased pseudotime trajectory showing the transition of MG into both activated and proliferative MGPC states (**a**). Expression *of RLBP1, GLUL, CDK1, TOP2A* were hallmarks of MG transition from the resting state (**a**). Heatmaps of MDK, PTN, CSPG5, and SDC4 were imposed on the pseudotime trajectory (**b,d**) and quantified for changes in gene expression on violin/scatter plots (**c**). Each dot represents a single cell, with black coloration indicative of two or more markers. MG from libraries of NMDA treated and NMDA treated with FGF2 plus insulin retinas were aggregated and projected onto a UMAP plot (**e**). These MG formed unique clusters categorized by their proliferative or activated gene expression profile. Heatmaps of *GLUL, RLBP1, VIM, TOP2A, PCNA,* and *CDK1* and violin plots of *GLUL, RLBP1, CDK1, TOP2A* demonstrate the differences in expression between retinal treatments (**e, g**) (**p<0.001, ***p<<0.0001; Wilcox rank sum with Bonferoni correction). The number on each violin indicates the percentage of expressing cells.

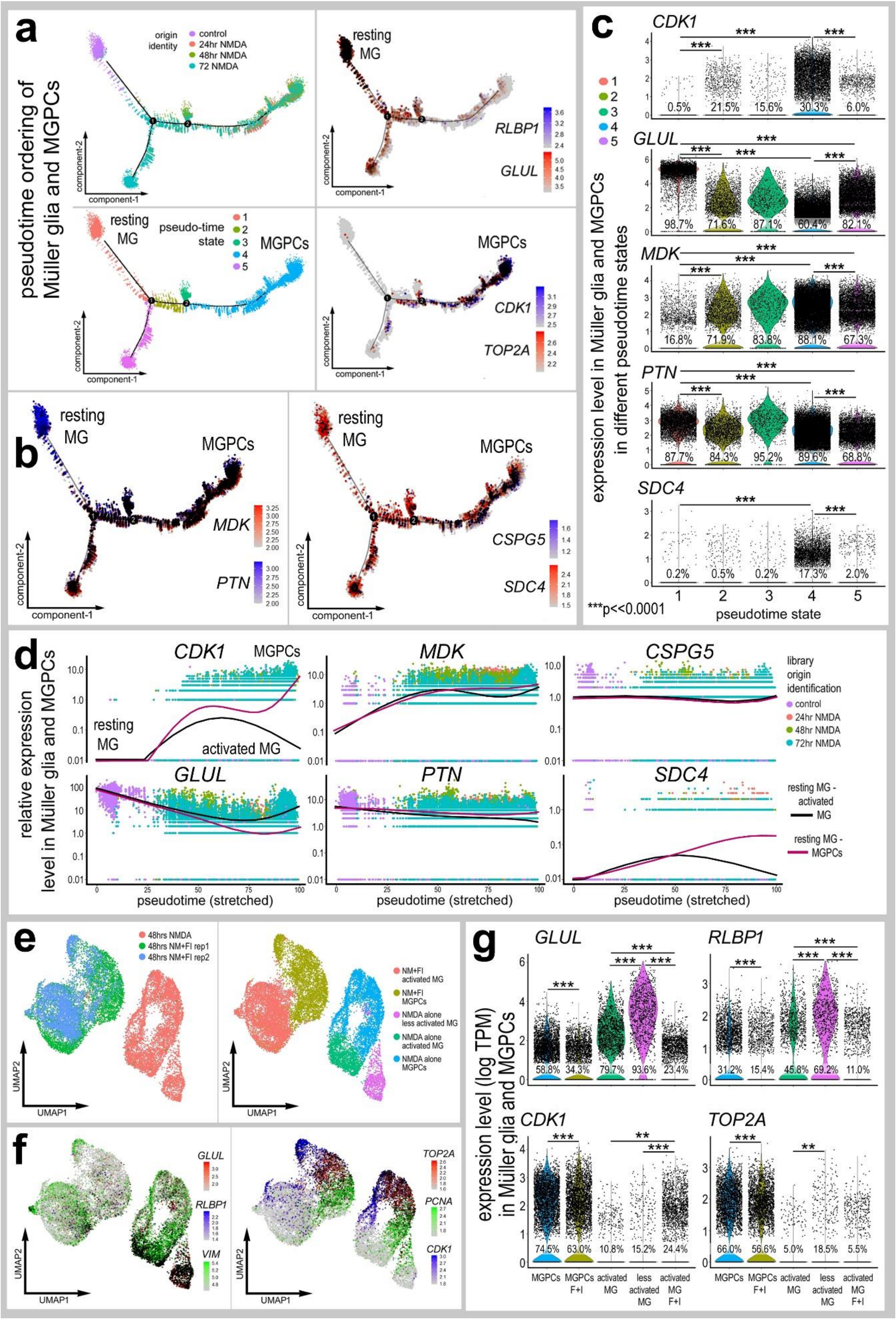
a
pseudotime ordering of Müller glia and MGPCs
origin identity
control
24hr NMDA
48hr NMDA
72 NMDA
resting MG
pseudo-time state
MGPCs
RLBP1
GLUL
CDK1
TOP2A
component-1
component-2
b
MDK
PTN
CSPG5
SDC4
c
CDK1
GLUL
MDK
PTN
SDC4
expression level in Müller glia and MGPCs in different pseudotime states
***p<<0.0001
pseudotime state
d
relative expression level in Müller glia and MGPCs
activated MG
pseudotime (stretched)
library origin identification
24hr NMDA
48hr NMDA
72hr NMDA
resting MG - activated MG
resting MG - MGPCs
e
48hrs NMDA
48hrs NM+FI rep1
48hrs NM+FI rep2
NM+FI activated MG
NM+FI MGPCs
NMDA alone less activated MG
NMDA alone activated MG
NMDA alone MGPCs
UMAP1
UMAP2
f
VIM
PCNA
g
expression level (log TPM) in Müller glia and MGPCs
MGPCs
MGPCs F+I
activated MG
less activated MG
activated MG F+I

**Supplemental Figure 4.** PAK1 inhibitor does not influence NIRG cells, microglia or numbers of dying cells. PAK1-specific inhibitor IPA3 was injected with and following NMDA (**a-d**) or before NMDA (**e-g**) and analyzed for accumulation and proliferation of NIRG cells, microglia and TUNEL+ cells. Alternatively, PP2A-specific inhibitors calyculin A or fostriecin were injected with and following NMDA and analyzed for accumulation and proliferation microglia or NIRG cells, and TUNEL+ cells (**h-v**). Sections of the retina were labeled for DAPI (blue), EdU (red) and CD45 (green; **h**), EdU (red) and Nkx2.2 (green; **o**), or TUNEL (red; **t**). Arrows indicate nuclei of proliferating microglia (**h**) or nuclei of NIRG cells (**o**). Arrow-heads indicate TUNEL-positive cells (**t**). The histogram/scatter-plots illustrate the mean (±SD) number of labeled cells. Each dot represents one biological replicate. Significance of difference (*p<0.05; ns – not significant) was determined by using a paired *t*-test.

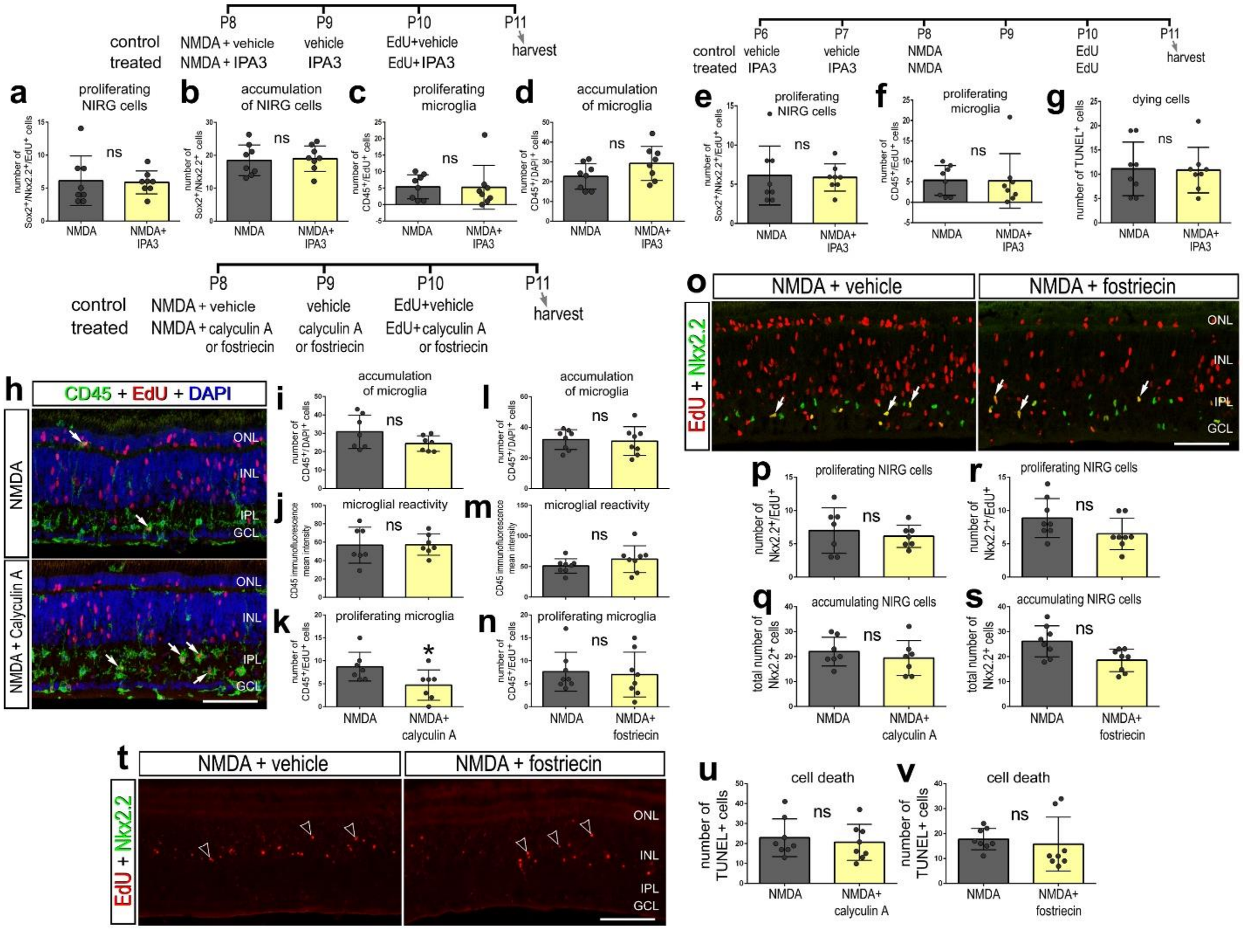

P8
P9
P10
P11
control
treated
NMDA + vehicle
NMDA + IPA3
vehicle
IPA3
EdU+vehicle
EdU+IPA3
harvest
P6
P7
P8
P9
P10
P11
vehicle
IPA3
NMDA
EdU
a
proliferating NIRG cells
b
accumulation of NIRG cells
c
proliferating microglia
d
accumulation of microglia
e
proliferating NIRG cells
f
proliferating microglia
g
dying cells
ns
NMDA
NMDA+ IPA3
NMDA + calyculin A or fostriecin
calyculin A or fostriecin
EdU+calyculin A or fostriecin
h
CD45 + EdU + DAPI
NMDA
NMDA + Calyculin A
ONL
INL
IPL
GCL
i
accumulation of microglia
j
microglial reactivity
k
proliferating microglia
l
accumulation of microglia
m
microglial reactivity
n
proliferating microglia
NMDA+ calyculin A
NMDA+ fostriecin
o
NMDA + vehicle
NMDA + fostriecin
EdU + Nkx2.2
p
proliferating NIRG cells
q
accumulating NIRG cells
r
proliferating NIRG cells
s
accumulating NIRG cells
t
u
cell death
v
cell death
number of TUNEL+ cells

**Supplemental Figure 5**. Expression of *MDK* and putative MDK-receptors in retinal cells treated with insulin and FGF2. scRNA-seq was used to identify patterns of expression of MDK-related genes among all types of retinal cells. In UMAP, violin and pseudotime plots, each dot represents one cell. Cells were obtained from control retinas (11,706 cells), and from retinas treated with 2 consecutive daily injections of insulin and FGF2 (9507 cells) or 3 consecutive daily injections of insulin and FGF2 (7237 cells) (**a**). UMAP plots revealed distinct clustering of different types of retinal cells; resting MG (4515 cells), 2 doses insulin and FGF2 MG (7685 cells), 3 doses insulin and FGF2 MG (3600 cells), MGPCs (2552 cells), NIRG cells (702 cells), oligodendrocytes (662 cells), bipolar cells (3182 cells), retinal ganglion cells (RGCs; 1159 cells), amacrine cells (2661 cells), rod photoreceptors (428 cells), and cone photoreceptors (1305 cells). MG were identified based on expression of *GLUL, RLBP1, VIM* and *SLC1A3* (**a**). MGPCs were identified based on expression of *NESTIN, CCNB2, CDK1* and *TOP2A* (**b**). UMAP feature plots revealed patterns of expression of *MDK, PTN, ITGB1, PAK1* and *CSPG5* (**c**). MG and MGPCs were re-embedded for unsupervised pseudotime ordering of cells which revealed trajectories with resting MG, proliferating MGPCs and activated MG from retinas treated with insulin+FGF2 largely confined to different branches in 3 different pseudotime states (**c, d**). Expression of *CDK1*, *GLUL, MDK, PTN, ITGB1, PAK1, and CSPG5* was projected onto pseudotime trajectories (**f**). Branched pseudotime plots and violin/scatter plots were generated to assess significant differences (***p<<0.001; Wilcox rank sum with Bonferoni correction) in *GLUL, MDK, PTN, PTN, ITGB1* and *PAK1* expression in different pseudotime states (**e**). The number on each violin indicates the percentage of expressing cells.

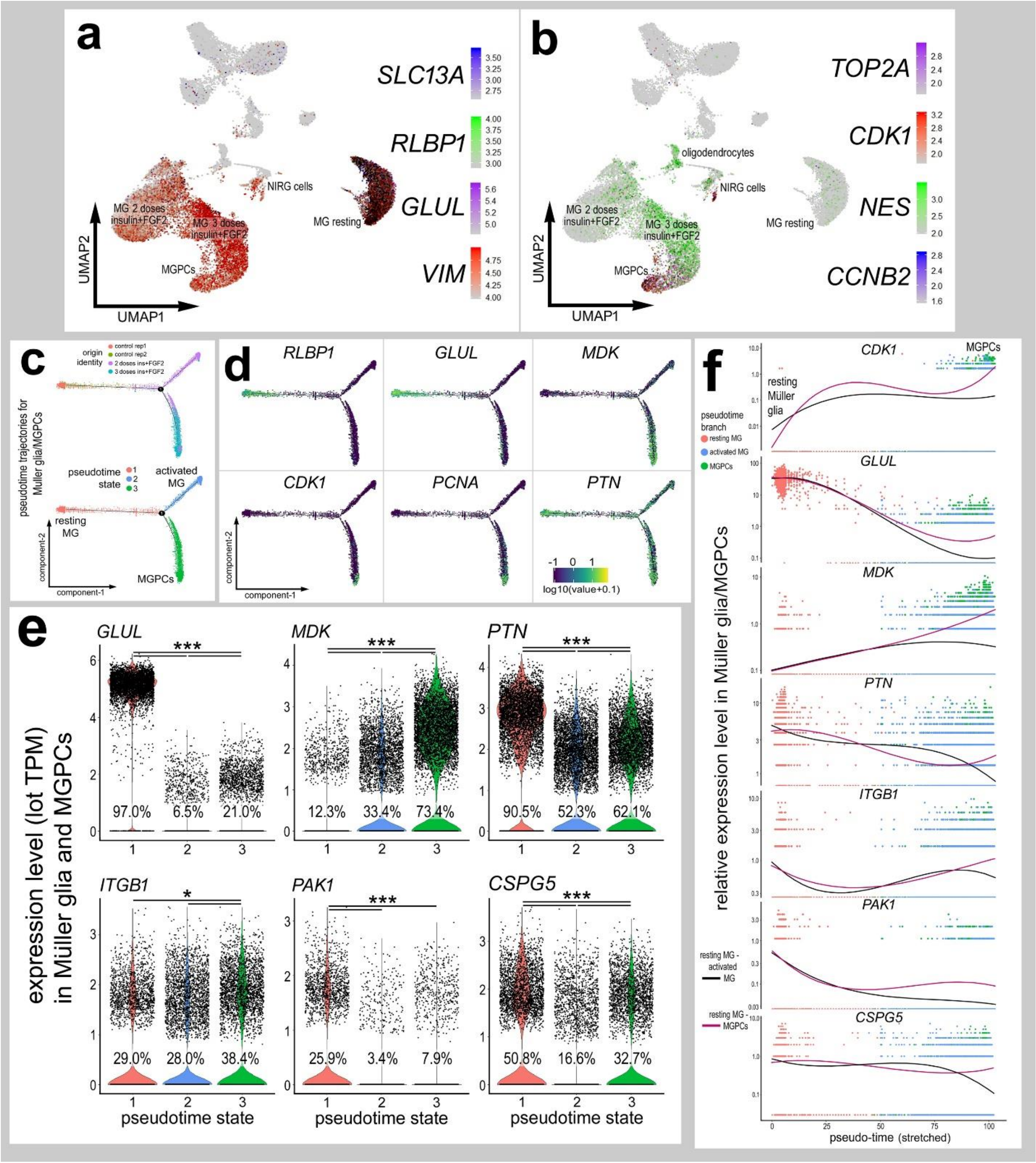
a
SLC13A
RLBP1
GLUL
VIM
NIRG cells
MG resting
MG 2 doses insulin+FGF2
MG 3 doses insulin+FGF2
MGPCs
UMAP2
UMAP1
b
TOP2A
CDK1
NES
CCNB2
oligodendrocytes
NIRG cells
MG resting
MG 2 doses insulin+FGF2
MG 3 doses insulin+FGF2
MGPCs
UMAP2
UMAP1
c
origin identity
control rep1
control rep2
2 doses ins+FGF2
3 doses ins+FGF2
pseudotime trajectories for Muller glia/MGPCs
pseudotime state
activated MG
resting MG
MGPCs
component-2
component-1
d
RLBP1
GLUL
MDK
CDK1
PCNA
PTN
log10(value+0.1)
e
expression level (lot TPM) in Müller glia and MGPCs
GLUL
MDK
PTN
ITGB1
PAK1
CSPG5
97.0%
6.5%
21.0%
12.3%
33.4%
73.4%
90.5%
52.3%
62.1%
29.0%
28.0%
38.4%
25.9%
3.4%
7.9%
50.8%
16.6%
32.7%
pseudotime state
f
relative expression level in Müller glia/MGPCs
CDK1
MGPCs
resting Müller glia
pseudotime branch
resting MG
activated MG
MGPCs
GLUL
MDK
PTN
ITGB1
PAK1
CSPG5
resting MG - activated MG
resting MG - MGPCs
pseudo-time (stretched)

**Supplemental Figure 6.** Expression Analysis of *Mdk*, *Ptn* and MDK-related genes in MG from normal, NMDA-damage, and FGF2 plus insulin treated mouse retinas. scRNA-seq was used to identify patterns of expression of MDK-related genes among acutely dissociated retinal cells (**a**). Unbiased pseudotime trajectories on MG clusters revealed a trajectory with resting MG to the left and activated MG from 3hrs after NMDA-treatment to the right with each dot representing a single MG cell from the aggregate library clustered into three individual states (**b, c**). The transition from resting to activated glia in the pseudotime trajectory can be visualized with a heatmap of *Glul, Scg2*, and *Srxn1* (d). Black coloration indicates cells with the expression of two or more markers. Heatmaps of *Pdk*, *Ptn*, *Cspg5*, *Itgb1*, and *Sdc4* demonstrate the changes in MDK-related markers in activated MG, and the transition of each gene is plotted in pseudotime (**e,f,g**). Libraries of retinas treated with NMDA and FGF2 and insulin were created and compared to NMDA treated libraries in UMAP and violin plots (**h, i**). In MG, *Glul, Vim, Ptn, Sdc4, Gfap, Mdk, Cspg5* and *Itgb1* genes were compared and analyzed for significant differences (*p<0.1, **p<0.0001, ***p<<0.0001; Wilcox rank sum with Bonferoni correction) (**j**). The number on each violin indicates the percentage of expressing cells.

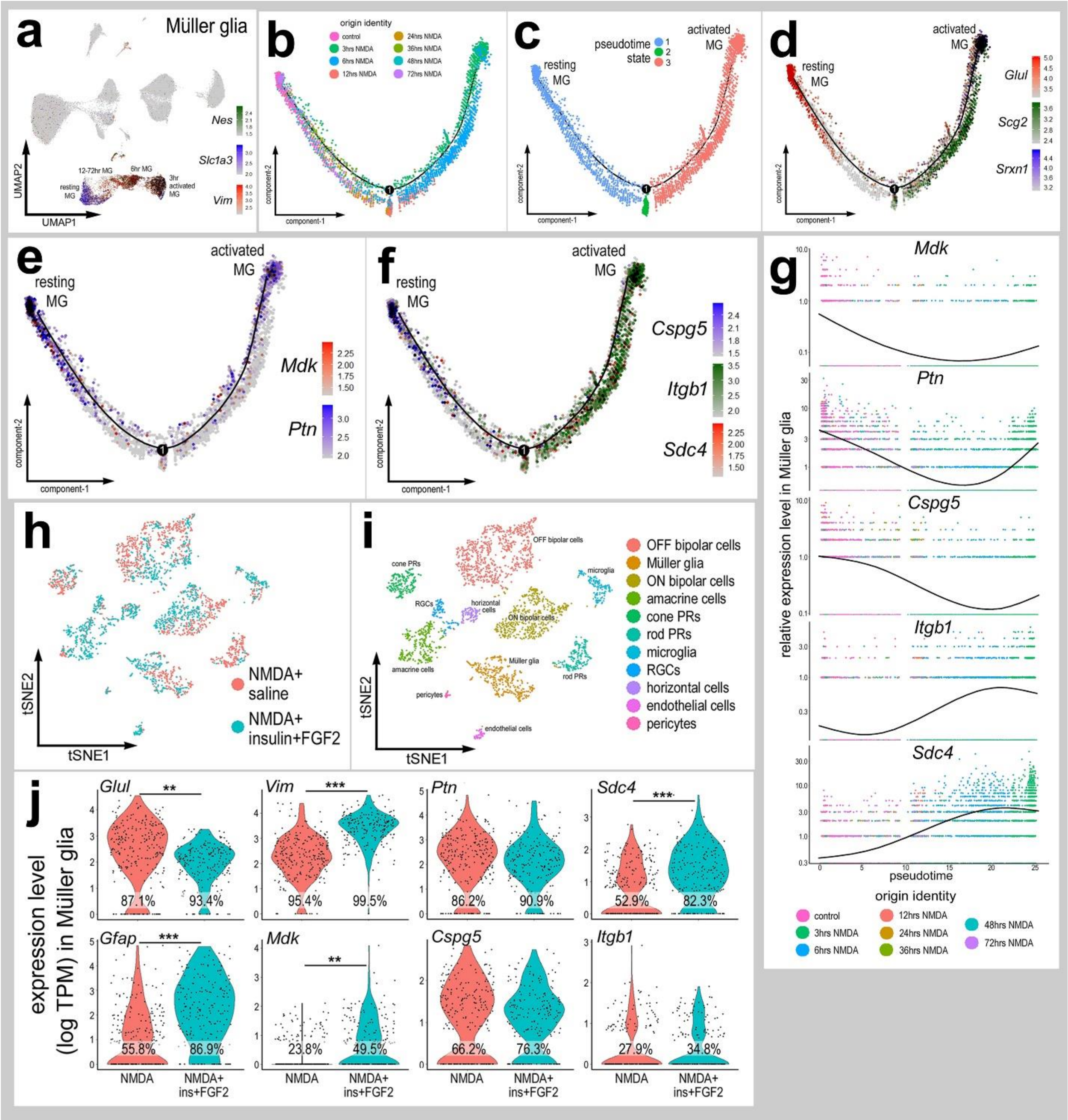
a
Müller glia
Nes
Slc1a3
Vim
UMAP1
UMAP2
resting MG
12-72hr MG
6hr MG
3hr activated MG
b
origin identity
control
3hrs NMDA
6hrs NMDA
12hrs NMDA
24hrs NMDA
36hrs NMDA
48hrs NMDA
72hrs NMDA
component-1
component-2
c
pseudotime state
resting MG
activated MG
d
Glul
Scg2
Srxn1
e
Mdk
Ptn
f
Cspg5
Itgb1
Sdc4
g
relative expression level in Müller glia
pseudotime
h
NMDA+ saline
NMDA+ insulin+FGF2
tSNE1
tSNE2
i
OFF bipolar cells
Müller glia
ON bipolar cells
amacrine cells
cone PRs
rod PRs
microglia
RGCs
horizontal cells
endothelial cells
pericytes
j
expression level (log TPM) in Müller glia
Glul
Vim
Ptn
Sdc4
Gfap
Mdk
Cspg5
Itgb1
87.1%
93.4%
95.4%
99.5%
86.2%
90.9%
52.9%
82.3%
55.8%
86.9%
23.8%
49.5%
66.2%
76.3%
27.9%
34.8%
NMDA
NMDA+ ins+FGF2
